# The interactome of the N-terminus of band 3 regulates red blood cell metabolism and storage quality

**DOI:** 10.1101/2020.11.30.404756

**Authors:** Aaron Issaian, Ariel Hay, Monika Dzieciatkowska, Domenico Roberti, Silverio Perrotta, Zsuzsanna Darula, Jasmina Redzic, Micheal P. Busch, Grier P. Page, Kirk C. Hansen, Elan Z Eisenmesser, James C Zimring, Angelo D’Alessandro

## Abstract

Band 3 (anion exchanger 1 - AE1) is the most abundant membrane protein in red blood cells (RBCs), the most abundant cell in the human body. A compelling model, based on indirect evidence, posits that - at high oxygen saturation - the N-term cytosolic domain of AE1 binds to and inhibits glycolytic enzymes, thus diverting metabolic fluxes to the pentose phosphate pathway to generate reducing equivalents. Dysfunction of this mechanism occurs during RBC aging or storage under blood bank conditions, suggesting a role for AE1 in the regulation of blood storage quality and efficacy of transfusion – a life-saving intervention for millions of recipients worldwide. Here we leverage two murine models carrying genetic ablations of AE1 to provide the first direct mechanistic evidence of its role in metabolic regulation and blood storage quality. Observations in mice phenocopied those in a human subject lacking expression of AE1_1-11_ (band 3 *Neapolis),* while common polymorphisms in the region coding for AE1_1-56_ increased susceptibility to osmotic hemolysis in healthy blood donors. Through thermal proteome profiling and cross-linking proteomics, we provide the first comprehensive analysis of the RBC interactome, with a focus on AE1_1-56_ and validate recombinant AE1 interactions with glyceraldehyde 3-phosphate dehydrogenase (GAPDH). Finally, we show that incubation with a cell-penetrating AE1_1-56_ peptide can rescue the metabolic defect in glutathione recycling and boost post-transfusion recoveries of stored RBCs from healthy human donors and genetically ablated mice, paving the way for the in vivo metabolic manipulation of RBCs facing oxidant stress – a landmark of many diseases.

**Key points:**

- Genetic ablation of N-term of band 3 results in significant metabolic aberrations and poor post-transfusion recoveries in mice and humans;
- Structural studies on the N-term of band 3 reveal a complex interactome with several enzymes, including GAPDH;

## Introduction

Band 3 - also known as anion exchanger 1 (AE1), owing to its role in the exchange of chloride for bicarbonate anions^1^-is one of the main targets of oxidant stress during red blood cell (RBC) aging in vivo and in the blood bank.^2,3^ AE1 also regulates RBC metabolism to allow adaptation to hypoxia and oxidant stress^4^. The N-terminal cytosolic domain of AE1 serves as a docking site for deoxygenated-hemoglobin (Hb), with significantly higher affinity than oxygenated-Hb^5,6^. Based on this evidence, it has been proposed that Hb, in addition to its central role of carrying O_2_ from the lungs to peripheral tissues, may serve as the “oxygen sensor” that directs AE1 to regulate metabolism appropriately. Indeed, studies had provided compelling, yet indirect evidence, that - when not encumbered by deoxy-Hb - the N-term of AE1 can bind to and, in so doing, inhibit the glycolytic enzyme glyceraldehyde 3-phosphate dehydrogenase (GAPDH).^7-11^ Notably, RBCs host ~1 million copies of both AE1 and GAPDH.^4^ A wealth of indirect evidence supports the likelihood and biological relevance of AE1-GAPDH interactions, based on data from immunofluorescence-based experiments,^7,8^ enzymatic activity assays, metabolic flux analysis via NMR,^9^ analytical ultracentrifugation^10^, in silico prediction,^11^ and genetic ablation of the N-term of band 3 in mice.^12^ However, early cross-linking studies have hitherto failed to produce direct evidence of the interaction of GAPDH with the N-term residues.^7^ Still, a widely accepted model has been proposed, according to which AE1 serves as a “railway switch” diverting glucose down different tracks (glycolysis vs. Pentose Phosphate Pathway – PPP) depending upon cellular needs^8,13,14^. This model explains why - when oxidant stress is high and GAPDH is bound to AE1 and thereby inhibited - RBC favor glucose oxidation via the PPP, to generate the reducing cofactor NADPH and fuel related antioxidant systems; ^15^ on the other hand, RBCs would rely on glycolysis when oxidant stress is low (e.g., high altitude^16^), Hb is deoxygenated and binds to the N-term of AE1, which in turn favors the release of glycolytic enzymes from the membrane to promote the generation of energy in the form of ATP and NADH. Regulating the balance between glycolysis and PPP is essential for RBCs to respond to their particular metabolic needs (membrane pump function, cytoskeleton and lipid homeostasis), which are a function of whether the RBCs are exposed to high or low oxygen tensions in the lungs or peripheral capillaries.^17^

We and others have proposed that this mechanism may be dysregulated as RBCs age and/or become damaged as a result of pathology or iatrogenic intervention (e.g., blood storage)^18,19^, depriving RBCs of critical metabolic plasticity and leading to their demise^3^. So far, the role of AE1 in RBC storage quality has only been hypothesized, based on a body of evidence that includes (i) an increase in AE1 fragmentation as a function of oxidant stress in stored RBCs;^20-24^ (ii) a progressive loss of the capacity to activate the PPP in stored RBCs,^25-27^ despite the gradual inhibition of GAPDH as a function of its oxidation at the active site Cys152 and 156, His179;^28^ (iii) an increase in band 3 phosphorylation at Y residues as a function of storage,^29^ which correlates to the release of microparticles from stored RBCs; (iv) disruption of GAPDH localization in the membrane upon phosphorylation of Y8 and 21 of AE1;^1,11,30^ (v) a poorer storage quality and post-transfusion recovery (PTR) of RBCs from donors who are incapable of activating the PPP because of deficient glucose 6-phosphate dehydrogenase (G6PD) activity;^31,32^ (vi) changes in AE1 oligomeric structure correlate with the loss of phospholipid asymmetry in stored RBCs, which could impact survival upon transfusion.^3,33^ Despite this evidence, the role of AE1 in blood storage quality has not yet been elucidated mechanistically.

## Methods

Methods are extensively detailed in the **Supplementary Methods extended**.

***Animal studies with mice*** All the animal studies described in this manuscript were reviewed and approved by the University of Virginia Institutional Animal Care and Use Committee (protocol n: 4269). Band 3 mouse founders, including huB3, HA Del and BS KO mice – originally generated by Chu et al.^34^ - were acquired from the National Institutes of Health mouse embryo repository and were bred with C57BL/6 females. The use of Ubi-GFP and HOD mice have been previously described in prior work from our group.^35^ Whole blood was drawn by cardiac puncture as a terminal procedure for the mice.

***Band 3 Neapolis*** RBCs (100 ul) were obtained from a subject carrying a mutation resulting in the lack of AE1 amino acids 1-11, as extensively described by Perrotta et al.^30^

***REDS-III RBC Omics study*** Details about patient enrollment, blood processing, osmotic hemolysis and development of the transfusion medicine genome wide genotyping array have been described extensively in prior studies on the background of the project.^32,36,37^

***Methylene blue treatments*** RBCs from WT and band 3 KO mice were incubated with methylene blue (100 uM, Sigma Aldrich) at 37°C for 1h, as described.^31^

***Storage under blood bank conditions*** RBCs from the four main strains investigated in this study, along with 13 different mouse strains described before^38^ were collected, processed, stored, transfused and post-transfusion recovery was determined as previously described.^38^

***Tracing experiments with labeled glucose, citrate, glutamine, arachidonic acid and methionine*** RBCs from all the mouse strains investigated in this study (100 ul) or RBC lysates from healthy donor volunteers (n=3) or the individual carrying band 3 Neapolis RBCs were incubated at 37°C for 1h in presence of stable isotope-labeled substrates, as described previously.^22^

***Metabolomics analyses*** were performed using a Vanquish UHPLC coupled online to a Q Exactive mass spectrometer (Thermo Fisher, Bremen, Germany), as described previously.^39,40^

***Proteomics*** Proteomics analyses, Thermal proteome profiling (TPP) and cross-linking experiments were performed via FASP digestion and nanoUHPLC-MS/MS identification (Thermo Fisher), as previously described.^41,42^ and extensively detailed in **Supplementary Methods extended**

***Isothermal Titration calorimetry (ITC)*** All ITC binding experiments were performed with a MicroCal iTC_200_ (Cytiva) set at 25 °C. GAPDH and Band 3 proteins were dialyzed into matching buffer (20 mM Bis-Tris pH 6.5, 50 mM NaCl, 2 mM DTT). The cell contained GAPDH at 0.1 mM while the syringe contained Band 3 at 1 mM. Reference power was set to 10 μcal/sec with a constant stirring speed of 1000 rpm. A total of 20 injections were performed. The first injection was excluded from data analysis. Experiments were performed in duplicate and the results were analyzed with the Origin ITC module.

***Nuclear Magnetic Resonance and structural models*** ^15^N-heteronuclear single quantum coherence (HSQC) spectra were collected at 25°C for the recombinantly expressed band 3 peptide 1-56 in the presence of 100 and 200 uM GAPDH. Data were collected on a Varian 900 using a standard ^15^N-HSQC sequence.

## Results

### Genetic ablation of the N-term of band 3 (AE1) impacts metabolism and glycolysis to pentose phosphate ratios in stored red blood cells

Despite similar function, the amino acid sequence is poorly conserved between mice and humans at the N-term^6^ (**Fig. 1.A**). As such, Low’s group generated a knock-in mouse by inserting the human binding site (amino acids (1-35)) into the mouse AE1, resulting in a human-mouse hybrid AE1 (HuB3) that could be used for a reductionist analysis of the human binding motif^5^. Two additional mouse models were subsequently made: (i) the first one, carries a deletion of the first 11 amino acid residues – hereon referred to as high affinity deletion or HA Del, since deletion of these residues negatively impacts binding of glycolytic enzymes, as suggested by immunofluorescence assays,^7,8^ enzymatic activity assays, metabolic flux analysis via NMR,^9^ analytical ultracentrifugation^10^, and in silico modeling;^11^ (ii) the second one was characterized by a deletion of amino acid residues 12-23, which resulted in the loss of affinity for the binding of deoxygenated-Hb (hereon referred to as binding site KO or BS KO)^34^ (**Fig. 1.B**). Based on this model (**Suppl. Fig.1.A-C**), here we hypothesized that HA Del and BS KO mice would store poorly, owing to the incapacity to activate the PPP as result of the lack of residues 1-11 or 12-23 of the N-terminus of AE1, respectively (**Fig.1. A-B**). Importantly, genetic ablation of AE1 N-term in these mice phenocopies the storage-induced fragmentation of AE1 N-term at multiple residues in human RBCs (**Suppl. Fig.1.D**). RBCs from WT, HuB3, HA Del and BS KO mice were stored under refrigerated conditions for 12 days, prior to metabolomics and unsupervised multivariate analyses (**Fig.1.C-D, Suppl.Table1**). The latter highlighted a significant impact of genotype on the end of storage levels of metabolites involved in glycolysis, the PPP and glutathione homeostasis (**Fig. 1.E; Suppl. Fig.2.A-B**). To further investigate the impact on glycolysis and the PPP, RBCs from these four mouse strains were incubated with 1,2,3-^13^C_3_-glucose upon stimulation with methylene blue (MB – **Figure 1.F**). MB is metabolized into leukomethylene blue in a reaction that consumes NADPH and thus promotes PPP activation by law of mass action.^31^ Determination of lactate isotopologues 2,3-^13^C_2_ and 1,2,3-^13^C_3_ – and their ratios – are indicative of the activation of the PPP and glycolysis, respectively, since the first carbon atom of 1,2,3-C3-glucose is lost in the form of CO_2_ in the oxidative phase of the PPP (**Fig.G-I, Suppl.Fig.2.C**). WT, HuB3 and BS KO mouse RBCs – but not HA Del - responded to MB stimulation by increasing fluxes through the PPP, a phenotype that was rescued (HA Del) or exacerbated (BS KO) upon incubation of RBC lysates with a recombinant AE1_1-56_ peptide (**Figure 1.I; Suppl.Fig.2.C**).

**Figure 1.**
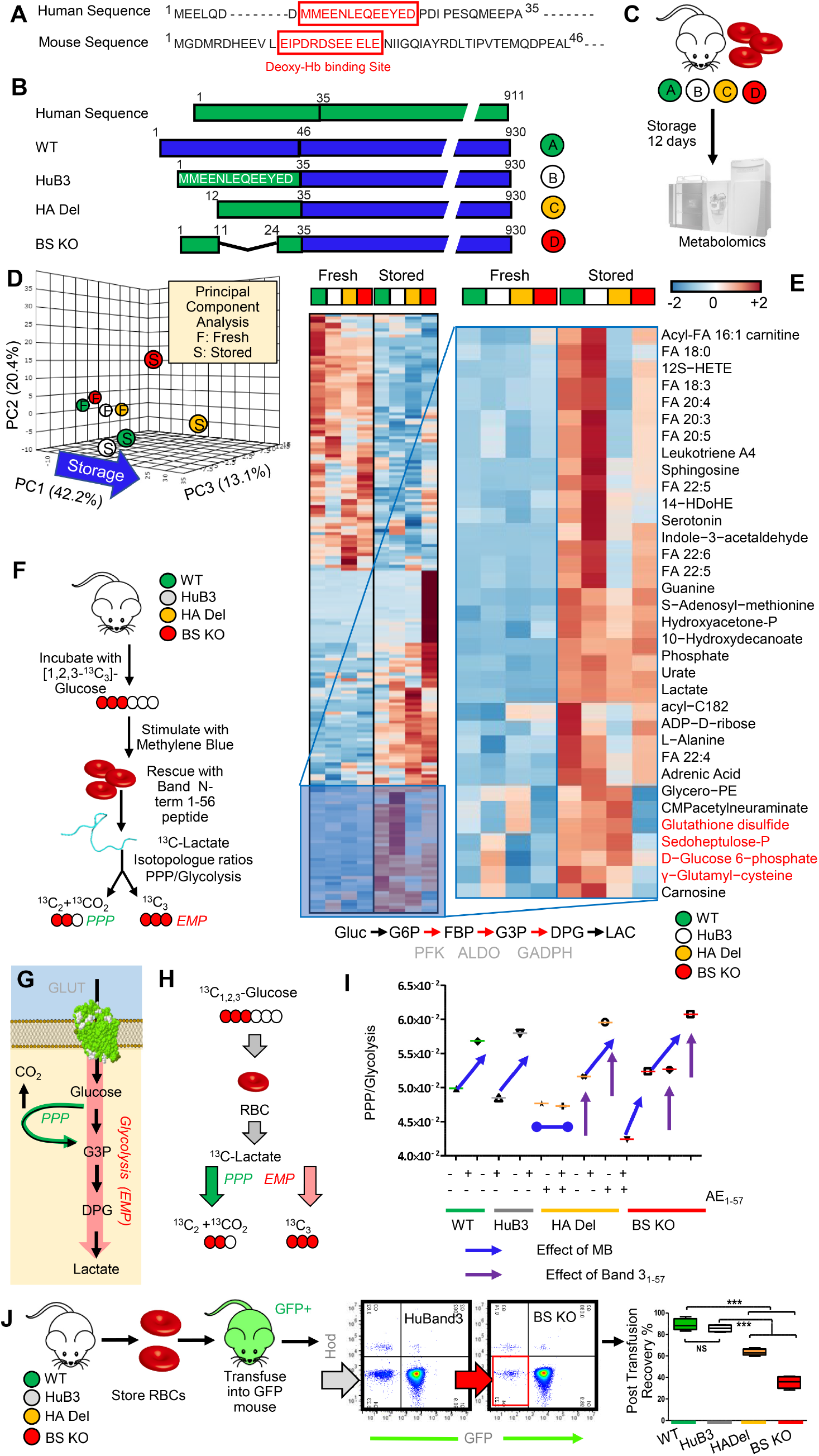
The N-term of band 3 (AE1) controls red blood cell metabolism during storage under blood bank conditions. Human and mouse N-term band 3 sequence are slightly divergent (**A**). To this end, in this study we leveraged humanized AE1 mice^12^, and compared them to WT mice (C57BL6), or humanized mice lacking residues 1-11 (deletion of the high affinity binding region for glycolytic enzyme, as previously suggested by immunofluorescence studies^12^ – HA Del) or 12-23 (hemoglobin binding site – BS KO) mice – **B**). RBCs from these mice were stored under refrigerated conditions for 12 days, prior to metabolomics analyses (**C**). Storage and genotypes had a significant impact on RBC metabolism, as gleaned by unsupervised principal component analysis (**D**) and hierarchical clustering analysis of significant metabolites by ANOVA (**E**). Specifically, strain-specific differences were noted in fresh and stored RBCs in glycolysis, pentose phosphate pathway (PPP) and glutathione homeostasis. To further investigate the impact on glycolysis and the PPP, RBCs from these four mouse strains were incubated with 1,2,3-C3-glucose upon stimulation with methylene blue (MB). Rescue experiments were also performed by incubating RBCs with a recombinantly expressed N-term AE1 peptide (residues 1-56 – **F**), prior to determination of the ratios of lactate isotopologues +3 and +2, deriving from glycolysis and the PPP, respectively (**G-H**). MB activates the PPP in all mouse strains except for the HA Del mice. Supplementation of the AE1_1-56_ peptide increases PPP activation and decreased glycolysis in HA Del mice and further exacerbated responses in all the other strains (**I**). Determination of end of storage PTR of RBCs from the four mouse strains showed significant decreases in PTR in the HA Del and BS KO mice (**J**).

### RBCs from mice lacking the N-term of band 3 have poorer post-transfusion recoveries

Fresh and stored RBCs from HA Del or BS KO mice also showed increased levels of carboxylic acids (succinate, fumarate, malate), acyl-carnitines (hexanoyl-, decanoyl-, dodecenoyl-, tetradecenoyl and hexadecanoyl-carnitine) and lipid peroxidation products compared to WT and HuB3 mouse RBCs, especially metabolites of the arachidonic and linoleic acid pathway (**Suppl.Fig.2.D-F**). Tracing experiments with labeled glutamine, citrate and methionine confirmed the steady-state aberrations in AE1 KO mice following MB challenge, while incubation with a recombinant AE1_1-56_ peptide only rescued glutaminolysis in HA Del mice and malate generation in BS KO mice (**Suppl.Fig.3.A-D**).

Expanding on previous targeted metabolomics studies,^35,38^ untargeted metabolomics analyses on fresh and stored RBCs from 13 different mouse strains identified metabolic correlates to PTR (**Suppl.Fig.2.G-J**), a Food and Drug Administration gold standard for storage quality. Pathway analyses and top ranked correlates are shown in **Suppl.Fig.2.F-G**, including previously reported oxylipins,^35,38^ but also purine oxidation products (xanthine), short chain and long chain saturated fatty acids, as well as carboylic acids as top negative correlates (**Suppl.Fig.2G-J, Supplementary Table 1**). All of these metabolites accumulated at higher levels in stored RBCs from the HA Del and BS KO mice compared to WT and HuB3 mice. Consistently, determination of end of storage PTR of RBCs from the four mouse strains showed significant decreases in PTR in the HA Del and BS KO mice (63.2 + 3.1% and 35.4 + 5.8%, respectively compared to 88.9 + 5.6% WT and 85.5 + 3.6% of HuB3 mouse RBCs, respectively – **Fig.1.J**).

### Lack of the 11 N-term amino acids of AE1 in humans (band 3 Neapolis) phenocopies RBC metabolism in AE1 KO mice

In 2005, Perrotta and colleagues^30^ identified a son of a consanguineous marriage with severe anemia, which was associated with a deficiency of AE1 expression (~10% of normal levels). Characterization of the gene coding for AE1 (gene name *SLC4A1)* revealed a single base substitution (T-->C) at position +2 in the donor splice site of intron 2, resulting in the generation of a novel mutant protein that lacked residues 1-11 (band 3 *Neapolis* – **Fig.2.A**). As such, the HA Del mice in this study *de facto* phenocopy band 3 *Neapolis* RBCs in humans. Consistent with observations in HA Del mice, omics analyses of band 3 *Neapolis* RBCs compared to controls showed significant alteration of glycolysis, glutathione homeostasis, decreases in methyl-group donors for isoaspartyl damage repairing^22^, increases in carboxylic acid, altered arginine and polyamine metabolism (**Fig.2.B**; **Suppl.Fig.4.A-B**). Incubation of band 3 *Neapolis* RBC lysates with 1,2,3-^13^C_3_-glucose confirmed a deficit in the capacity to activate the PPP following stimulation with MB, a defect that was corrected by rescue with a recombinant AE1_1-56_ peptide (**Fig.2.C**), similar to HA Del RBCs.

**Figure 2.**
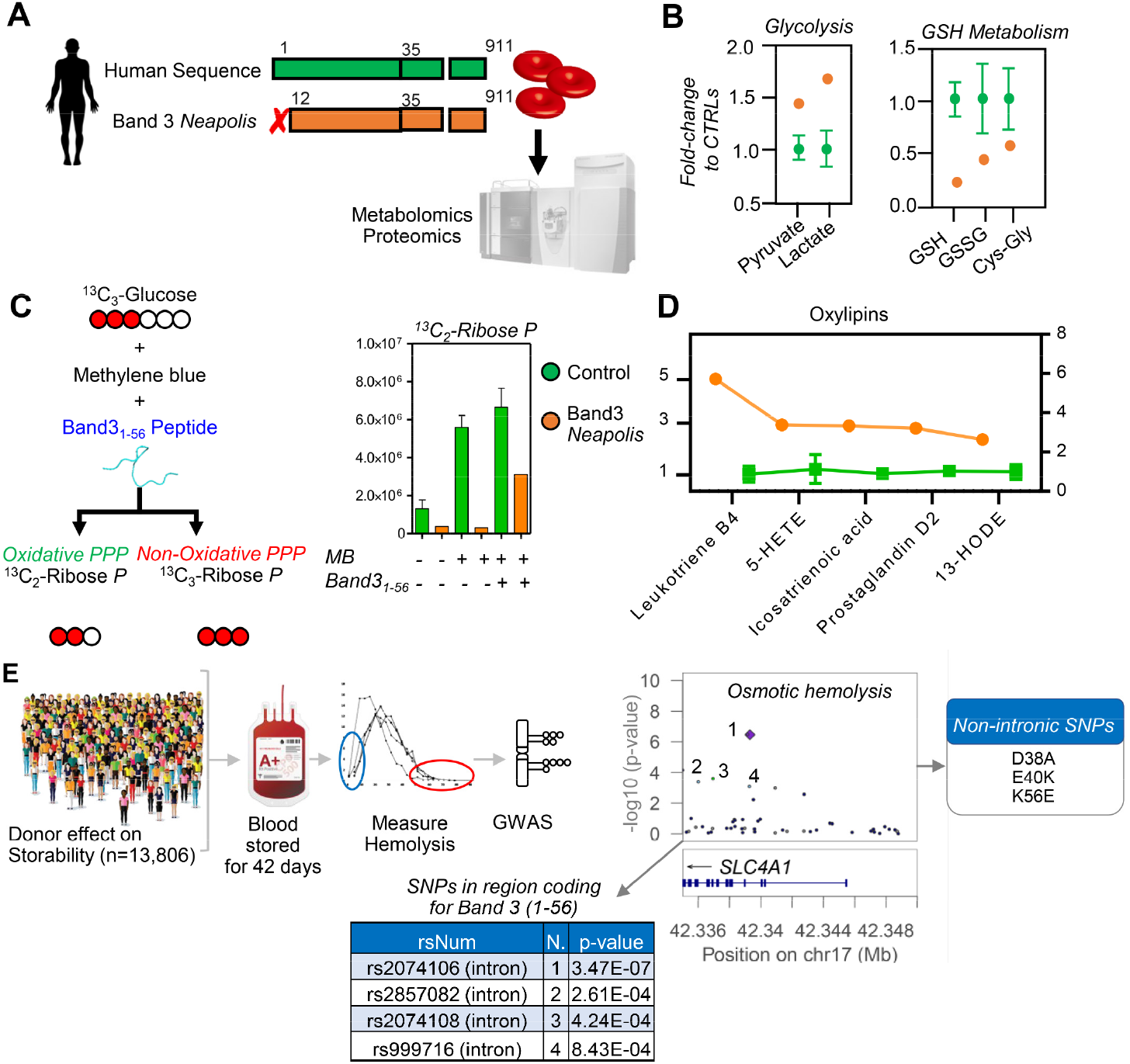
Metabolic impact of AE1 1-11 deletion in human RBCs in band 3 Neapolis and alterations of osmotic fragility of stored RBCs from donors with polymorphisms in AE1_1-56_. Lack of amino acid residues 1-11 in humans (band 3 *Neapolis-* **A**) results in increased activation of glycolysis and decreases in glutathione pools (**B**). Tracing experiments with 1,2,3-^13^C3-glucose show impaired responses to methylene blue (MB)-induced activation of the pentose phosphate pathway (PPP) compared to glycolysis in band 3 Neapolis, a phenotype that is partially rescued in vitro by supplementation of a recombinant AE1_1-56_ peptide (**C**). de facto phenocopying HA Del mice in this study. However, band 3 Neapolis RBCs were also characterized by higher levels of oxylipins (**D**), comparable to those observed in stored BS KO mice. GWAS studies on 13,806 healthy donor volunteers revealed an increased in osmotic fragility in subjects carrying polymorphism in the region coding for band 3 (gene name *SLC4A1)* residues 1-56 and neighboring introns (**E**).

Proteomics analyses showed higher levels of hexokinase, but lower levels of all the remaining glycolytic enzymes downstream to it, as well as of enzymes of the PPP and glutathione synthesis in the RBCs from the band 3 *Neapolis* individual (**Suppl.Fig.4.C**), despite comparable mean corpuscular value, mean corpuscular hemoglobin and hematocrit (**Suppl.Table1**). Alterations in RBC metabolism in band 3 *Neapolis* RBCs comparable to those observed in AE1 KO mice were confirmed not just at steady state (e.g., higher levels of lipid oxidation products – **Fig.2.D**), but also by tracing experiments with ^13^C ^15^N-glutamine; ^13^C-citrate, ^13^C ^15^N-methionine, and deuterium-labeled arachidonate (**Suppl.Fig.4.D-G**).

### Single Nucleotide Polymorphisms (SNPs) in the region coding for AE1_1-56_ correlate with increased osmotic fragility of stored human RBCs

A large cohort of 13,806 healthy donor volunteers was enrolled in the Recipient Epidemiology and Donor assessment Study – REDS III-RBC Omics.^32^ Units from these donors were stored for up to 42 days prior to determination of the RBC susceptibility to hemolysis following osmotic insults.^37^ Genome wide association studies (GWAS) of over 879,000 SNPs^36^ revealed that the highly polymorphic region on chromosome 17 coding for AE1 (gene name *SLC4A1)* ranked amongst the top correlates to osmotic fragility in the whole study (**Fig.2.E**).

### Proteomics and structural studies on the interactome of band 3

After confirming the lack of the N-term portion of AE1 in the KO mice (**Suppl.Fig.3.E-G**), we characterized the proteomes of RBCs from the four mouse strains, with a focus on protein levels that were similarly altered in both HA Del or BS KO mice, or uniquely altered in either KO (**Suppl.Fig.3.H-K**). The list of proteins whose levels were decreased in band 3 KO mice included numerous proposed interactors of AE1 (in red in **Suppl.Fig.3.H-K, Suppl.Table1**). This observation could be explained either by altered protein expression in erythroid cells during maturation to RBCs, or could be indicative of a protective role from proteolytic degradation by interaction of these proteins with the N-term of AE1, as proposed in other studies of the RBC degradome.^43^

To test the latter hypothesis, we used thermal proteome profiling (TPP) coupled to tandem mass tag 10 (TMT10 – **Fig.3.A**) to determine candidate interacting partners to AE1_1-56_. Candidate interactors are identified as those with the most extreme alterations in the temperature at which their solubility decreases/increases and precipitation is observed (ΔTm – **Fig.3.B, Suppl.Table1**). A few representative melting curves for the top hits for proteins stabilized or destabilized by the presence of AE1_1-56_ (red) compared to untreated controls (blue) is provided in **Fig.3.C.** Hits included several glycolytic enzymes GAPDH, aldolase A (ALDOA), phosphoglycerate kinase (PGK) and lactate dehydrogenase A (LDHA); hemoglobin alpha and beta (HBA and HBB); enzymes involved in redox homeostasis and oxidant damage-repair (protein L-isoaspartate O-methyl-transferase (PIMT), peroxiredoxin 2 (PRDx2), gamma-glutamyl cysteine ligase (GCLC), glyoxalase 1 (GLO1) aldehyde dehydrogenase 1 (ALDH1A1), glutathione peroxidase 4 (GPX4), phosphogluconate dehydrogenase (PGD)) or other metabolic enzymes (acetyl-CoA lyase (ACLY), arginase 1, adenosyl homocysteine hydrolase (AHCY), glutamate oxalacetate transaminase (GOT1), adenylate kinase (ADK) – **Figure 3.C**.

**Figure 3.**
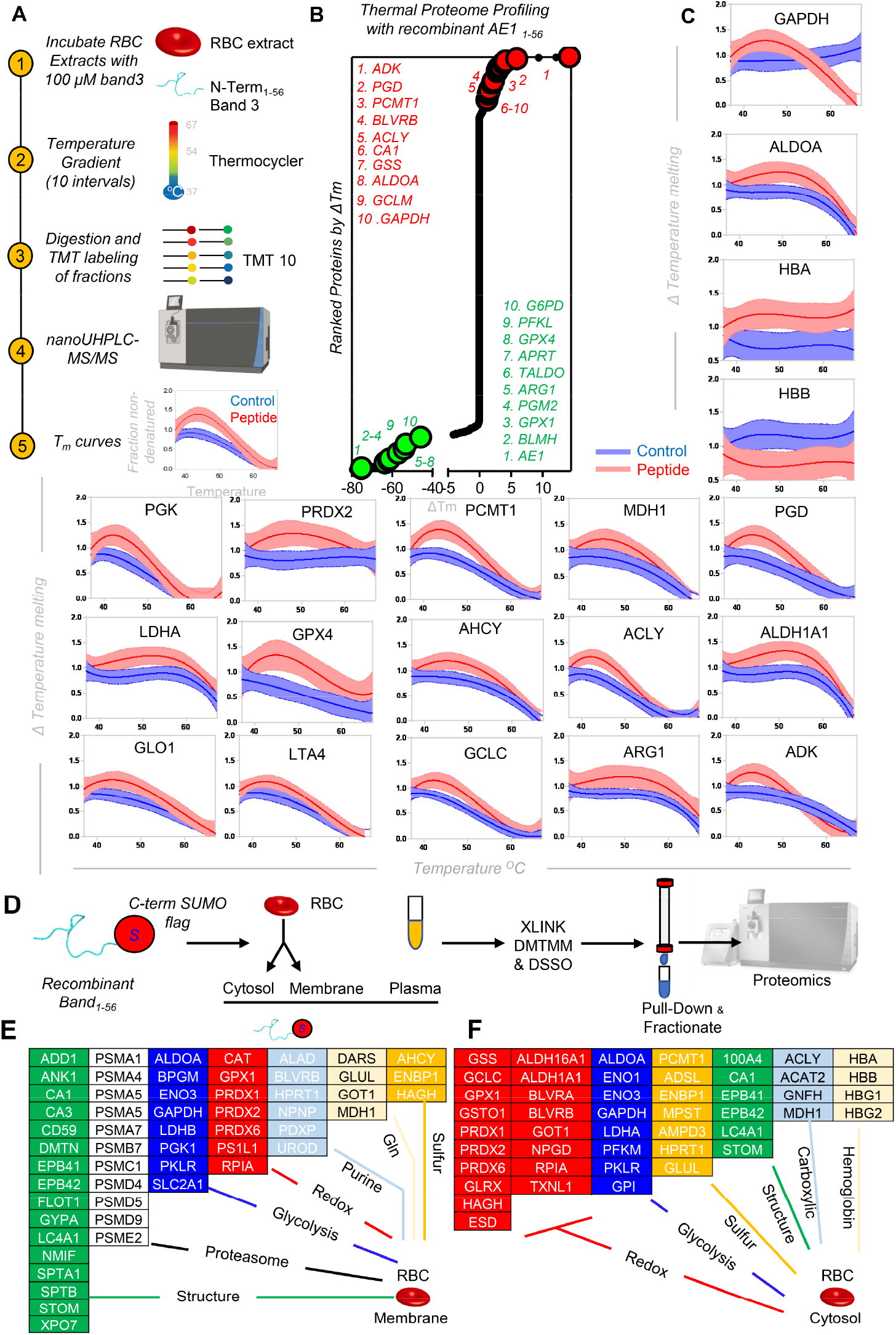
Thermal proteome profiling and Xlinking proteomics of recombinant peptide 1-56 of band 3 in RBC lysates. Thermal proteome profiling (TPP) experiments were performed by incubating RBC lysates with a recombinantly expressed peptide coding for the amino acids 1-56 in the N-term of band 3 at a gradient of temperature from 37-67°C, labeling with 10 different tandem mass tags (TMT 10), pooling and analysis via nanoUHPLC-MS/MS (**A**). Proteins were thus ranked as a function of their alterations in the temperature at which their solubility decreases and precipitation is observed (ΔTm – **B**). A few representative melting curves for the top hits for proteins stabilized or destabilized by the presence of the band 3 peptide 1-56 (red) compared to untreated controls (blue) is provided in **C**. A peptide coding for the amino acids 1-56 of the N-term of band 3 was recombinantly expressed with a SUMO-tag and/or a His-Flag tag at either the N- or C-term terminus of the peptide, prior to incubation with plasma, red blood cell cytosols and membrane in independent experiments, enrichment in Nichel columns, pull-down against the SUMO tag, and cross-linking with agents disuccinimidyl sulfoxide (DSSO) or 4-(4,6-dimetoxy-1,3,5-triazin-2-yl)-4-metylmorpholinium – DMTMM), prior to protein digestion, fractionation of cross-linked peptides and nanoUHPLC-MS/MS-based identification of band 3 interacting partners via MS2/MS3 analyses (**D**). Top interactors for RBC membrane and cytosol interactors are listed in **E-F**, respectively, divided by pathway.

### Immuno-precipitation and cross-linking proteomics studies with recombinantly expressed AE1_1-56_

To confirm and expand on the TPP-TMT10 analyses, we recombinantly expressed AE1_1-56_ with a His-SUMO-tag at either the N- or C-terminus, prior to incubation with human plasma, RBC cytosols and membrane from human RBC lysates and purification/pull-down of interacting partners **Suppl.Fig.5.A**). Pathway analyses of the hits from these analyses confirmed most of the hits from TPP (**Suppl.Table1**), revealing a widespread network of interactors of AE1, including up to 63 proteins involved in metabolic regulation, as mapped against the KEGG pathway map of human metabolism (**Suppl.Fig.5.B-C**). One limitation of the TPP and immunoprecipitation approaches is that they identify both direct and indirect interactors to AE1 that can be pulled down because of their complexing with direct AE1 interactors. To disambiguate between direct and indirect interactors to AE1_1-56_, we repeated the precipitation experiments by also introducing cross-linking agents disuccinimidyl sulfoxide (DSSO) or 4-(4,6-dimetoxy-1,3,5-triazin-2-yl)-4-metylmorpholinium – DMTMM – **Fig.4.A-B, Suppl.Table1**). As proof of principle, we first performed these analyses on RBC lysates without the addition of recombinant AE1, providing the first experimental report of the RBC interactome (shown in the form of circos plot or network view in **Fig.4.C-D**, respectively; **Suppl.Table1)**, significantly expanding on *in silico* predictions in the literature.^44^ This analysis allowed to recapitulate decades of structural studies on RBC hemoglobin (**Suppl.Fig.6**) and the RBC membrane interactome (**Fig.5.A-C**), relevant to the regulation of RBC cytoskeleton and the role AE1 plays as a lynchpin for RBC membrane structural proteins (**Fig.5.C**), AE1 multimerization in the intra and extra-cellular compartment (**Fig.5.D-E**).

**Figure 4.**
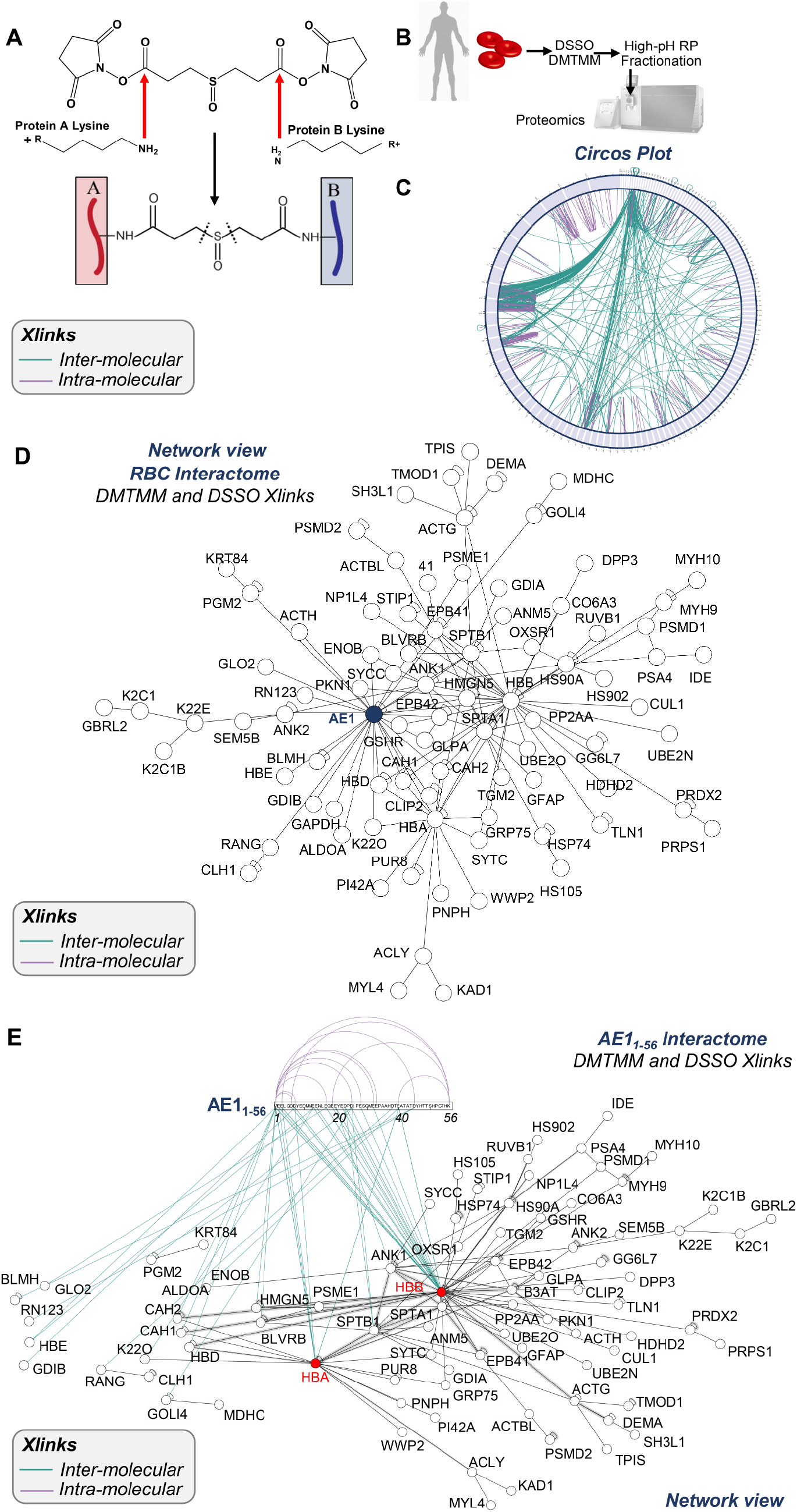
Experimental RBC interactome, as gleaned by cross-linking mass spectrometry. Focus on the interactome of AE1_1-56_. In **A**, an overview of the cross-linking strategy with DSSO. This approach affords determination of protein-protein interactions by cross-linking free amines of adjacent residues of interacting proteins within a range spanning from 11.4 to 24 Å. In the present study, DSSO and DMTMM (E/D residues to K) were combined to investigate the RBC interactome (**B**). In **C**, circos plot view of the protein-protein cross-links identified in this study (full list in **Supplementary Table 1**). In **D**, the same inputs were used to provide a network view of the results. In **E**, an interactomics overview merging the data from all the cross-linking proteomics studies, showing direct and indirect interactors with band 3 1-56 and the residues on band 3 with which the proteins were identified to cross-link to.

**Figure 5.**
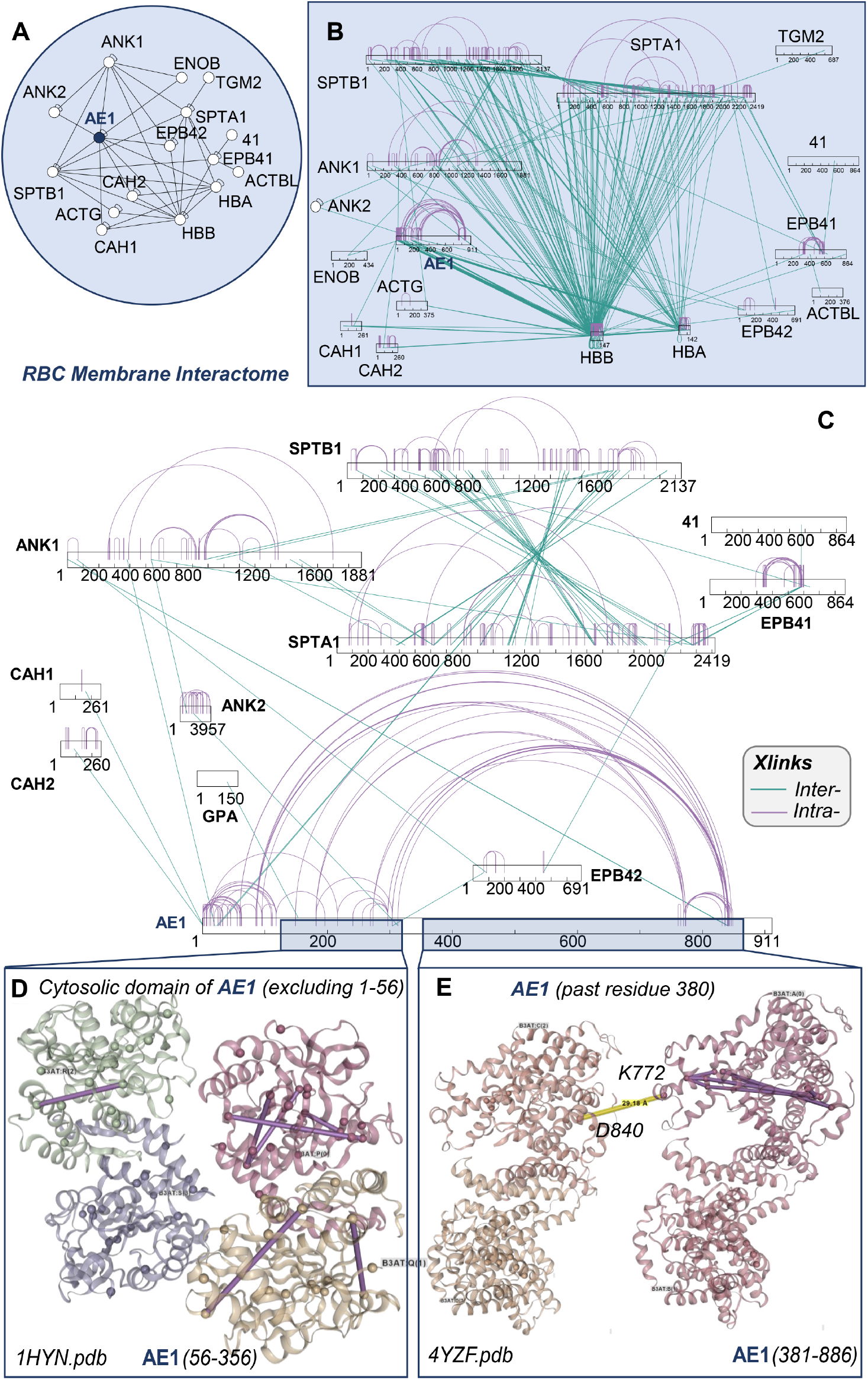
Experimental interactome of red blood cell membrane proteins. In **A**, highlighted network of RBC membrane proteins according to the canonical models (proteins are abbreviated as Uniprot gene names). In **B**, an overview of the interactions across these proteins in bar plot format. In **C**, expansion of the bar plots focusing on band 3 (B3AT), spectrin alpha and beta (SPTA1 and SPTB1), ankyrins 1 and 2 (ANK1 and ANK2), carbonic anhydrase 1 and 2 (CAH1 and CAH2), erythrocyte protein band 4.1 and 4.2 (41 and EPB42). In **D** and **E**, highlights of the intramolecular interaction of band 3 (residues 56-356 – *1HYN.pdb* and 381-886 – *4YZF.pdb,* respectively). In **F**, zoom in on a sub-network of structural proteins interacting with STPB1. Zoom in on a subset of intramolecular cross-links in SPTB1 (*1S35.pdb* – **G**) are shown in **H**.

To further focus on the AE1 N-term, crosslinking proteomics studies were repeated by adding recombinant AE1_1-56_. This approach afforded determination of direct AE1_1-56_ protein interactors by identifying cross-linked free epsilon-amines of K with adjacent identical residues (DSSO) or carbon atoms in the carboxylic groups of D or E amino acids (DMTMM – **Fig.4.A**), provided these residues face each other within a range spanning from 11.4 to 24 Å.^45^ The list (**Suppl.Table1)** included several structural proteins, proteasomal components, and enzymes involved in glycolysis, redox homeostasis, purine, glutamine or sulfur metabolism – both in RBC membranes and cytosols (**Fig.3.E-F**, respectively). An overview of the AE1_1-56_ interactome is shown in **Fig.4.E**. This approach revealed three hot spots of cross-links, at D or E residues in amino acid sequences 1-7, 13-25 and 40-45 of AE1 N-term (**Fig.4.E**). Intramolecular and inter-molecular interactions were observed for AE1 itself at the N-term (positions 45-6; 45-38; 45-56; 45-1; 25-1, 38-1 etc.) and C-term (833-840; 837-765; 771-849, etc. – **Suppl.Table1**). We report direct interactions of AE1_1-56_ with hemoglobin beta (HBB – 1-2; 25-2; 23-2; 10-2; 10-134; 1-97; 38-2; 6-2; 1-100 – **Suppl.Fig.7**) and hemoglobin alpha (HBA – 38-7, 38-86; 1-97 less frequent and with lower scores than the cross-links with HBB).^5^ In addition, direct cross-links were identified for AE1_1-56_ with spectrin beta (SPTB1 – residues 23 to 1664, respectively), ankyrin (residue 10 – 381 – **Fig.5.A-C**), phosphoglucomutase 2 (residues 23-88), enolase beta (38-198). Glyoxylase 2 (10-299); Rab GDP dissociation inhibitor beta (position 2-2); (**Suppl.Table1**), Golgi integral membrane protein 4 (23-570 – consistent with work on endosomes);^46^ carbonic anhydrase 2 (1-81); biliverdin reductase B (1-40).

Pull-down of AE1_1-56_ followed by cross-linking studies identified further candidate indirect interactors that are either part of macromolecular complexes with direct AE1_1-56_ interactors, including peroxiredoxin 2,^47^ peroxiredoxin 6, catalase, phosphofructokinase isoforms,^12^ protein 4.1 and 4.2, glucose phosphate isomerase, stomatin, adducin beta, carbonic anhydrase 1, 1-phosphatidylinositol 4,5-bisphosphate phosphodiesterase eta-2 (full list in **Suppl.Table1**). Other enzymes were pulled-down, but not directly cross-linked (suggestive of an indirect interaction with AE1, though direct interactions with other residues beyond residue 56 cannot be ruled out), such as ACLY, bleomycin hydrolase, Serine/threonine-protein kinase OSR1, desmoplakin, calpastatin, glycophorin A.

### Biophysical studies of the GAPDH and AE1_1-56_ interaction

The interaction between recombinant GAPDH tetramers (**Fig.6.A**) and AE1_1-56_ (also including K56) was confirmed via ITC with a Kd of 2.56 uM (**Fig.6.B**), and 2D-NMR upon titration of GAPDH (0, 100 and 200 uM – **Fig.6.C**). Of note, recombinant expression of an AE1 peptide spanning from residues 1-30 resulted in weaker interactions with GAPDH (Kd: 44.8 uM – **Suppl.Fig.8.A-B**). Calculations of chemical shift perturbations (CSPs) induced by GAPDH titration to the 15N-labeled AE1_1-56_ (**Fig.6.D**) provided clues on the structural organization of the otherwise intrinsically disordered N-term of AE1. Specifically, residues that comprise the AE1 binding site exhibit severe line-broadening due to exchange with the much larger GAPDH. For example, the terminal residues are the primary binding site as illustrated by the near complete disappearance of the central Y8 amide, compared to the amide of Y48 that exhibits no CSP (**Fig.6.E**). DSSO and DMTMM cross-links of GAPDH and AE1_1-56_ (a representative spectrum between AE1 K56 to GAPDH D203 is shown in **Fig.6.F**) are reported in **Fig.6.G** (**Supplementary Table 1**). Cross-linked residues are mapped against GAPDH monomeric structure in **Fig.6.H**. Similar results were obtained when probing interactions between GAPDH and AE1_1-30_ (**Suppl.Fig.8.C-E**), with the exception of cross-links between E40, a hot spot for interaction with several GAPDH residues in the longer AE1 peptide (3, 5, 84, 86, 194, 219, 251, 259, 260, 263 – **Fig.6.G**). These experiments provide further proof of a direct interaction of the N-terminus of AE1 to GAPDH and indicate that the N-terminal region comprising residues 1-20 is the primary binding site with additional contacts that also contribute to the AE1/GAPDH interaction. ITC, NMR and cross-linking proteomics data were used to build a model of AE1_1-56_ interaction with GAPDH with the software Rosetta, resulting in the active site pocket of GAPDH being exposed to residues 1-15 of AE1 (**Fig.6.I-K**; distances between K and D/E residues are shown in **Suppl.Fig.8.F**).

**Figure 6.**
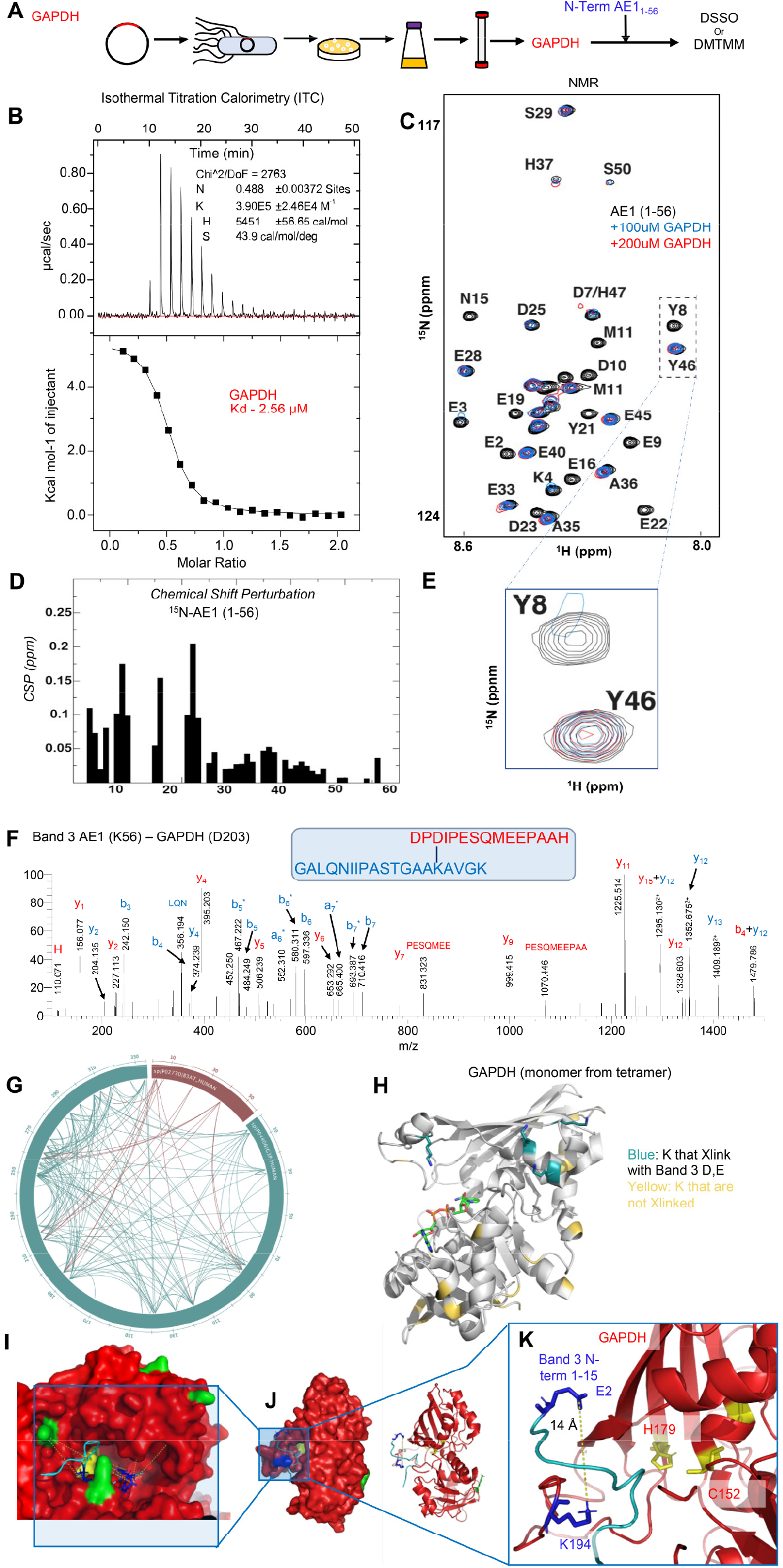
Structural studies of recombinantly expressed glyceraldehyde 3-phosphate dehydrogenase (GAPDH) and its interaction with the N-term of band 3 (residues 1-56). GAPDH was recombinantly expressed in *E.coli* prior to purification and interaction studies with the recombinantly expressed band 3 peptide (residues 1-56 – **A**). These studies included isothermal titration calorimetry (**B**), nucelar magnetic resonance of band 3 (1-56) with 100 or 200 uM GAPDH (**C**) and derived calculation of chemical shift perturbations (CSP – **D**), in silico modeling of band 3 (1-56) structure based on NMR data (**E**). Following these studies, cross-linking proteomics analyses were performed in vitro by co-incubating GAPDH and band 3 (1-56) in presence of DSSO or DMTMM. In **F**, a representative spectrum from one of the most abundant cross-links, which are comprehensively mapped in the circos plot (light blue for intramolecular cross-links, red for intermolecular ones) in **G**. Results were thus mapped against GAPDH monomeric structure in **H** (blue: all lysine residue on GAPDH that were experimentally found to cross-link to band 3 aspartyl or glutamyl side chains, vs in yellow the lysine that were available for crosslinking but were not found to face band 3 acidic residues within the reach of the crosslinker of ~20 Å). In **H-J-K**, NMR and cross-linking proteomics data were used to build a model of band 3 (1-56) interaction with GAPDH with the software Rosetta, resulting in the active site pocket of GAPDH being exposed to residues 1-15 of band 3.

### Cell membrane-permeable AE1_1-56_ peptides reverse the defect in the glutathione recycling capacity of RBCs from band 3 KO mice

To rescue the genetic defect resulting from ablation of the AE1 N-term, we designed three membrane permeable versions^48^ of AE1_1-56_ through addition of a polyarginine (polyArg), internalization sequence or TAT sequence at the C-term of the peptide (**Fig.7.A**). Human RBCs were thus incubated with a control (scramble), an AE1_1-56_ peptide (non-penetrating) or the three penetrating peptides in presence of 1,2,3-^13^C_3_-glucose and methylene blue stimulation to activate the PPP (**Fig.7.A**). Tracing experiments show that PPP activated following MB in all cases, but ATP synthesis was only preserved in the RBCs treated with the cell penetrating peptides (**Fig.7.B**), consistent with the schematic in **Fig.7.C**. We thus hypothesized that incubation with polyArg-AE1_1-56_ would at least partially rescue the defect in PPP activation upon MB stimulation of RBCs from AE1 KO mice, especially HA Del (**Fig.7.D**). Results show increased glutathione oxidation (GSSG/GSH ratios) in response to MB treatment in all cases, but significant rescue by the AE1 peptide in both AE1 KOs. Storage of murine RBCs from the four mouse strains in presence of polyArg-AE1_1-56_ promoted increases in PTR in WT and HA Del mice back to levels observed in HuB3, but not in BS KO mice (**Fig.7.E**).

**Figure 7.**
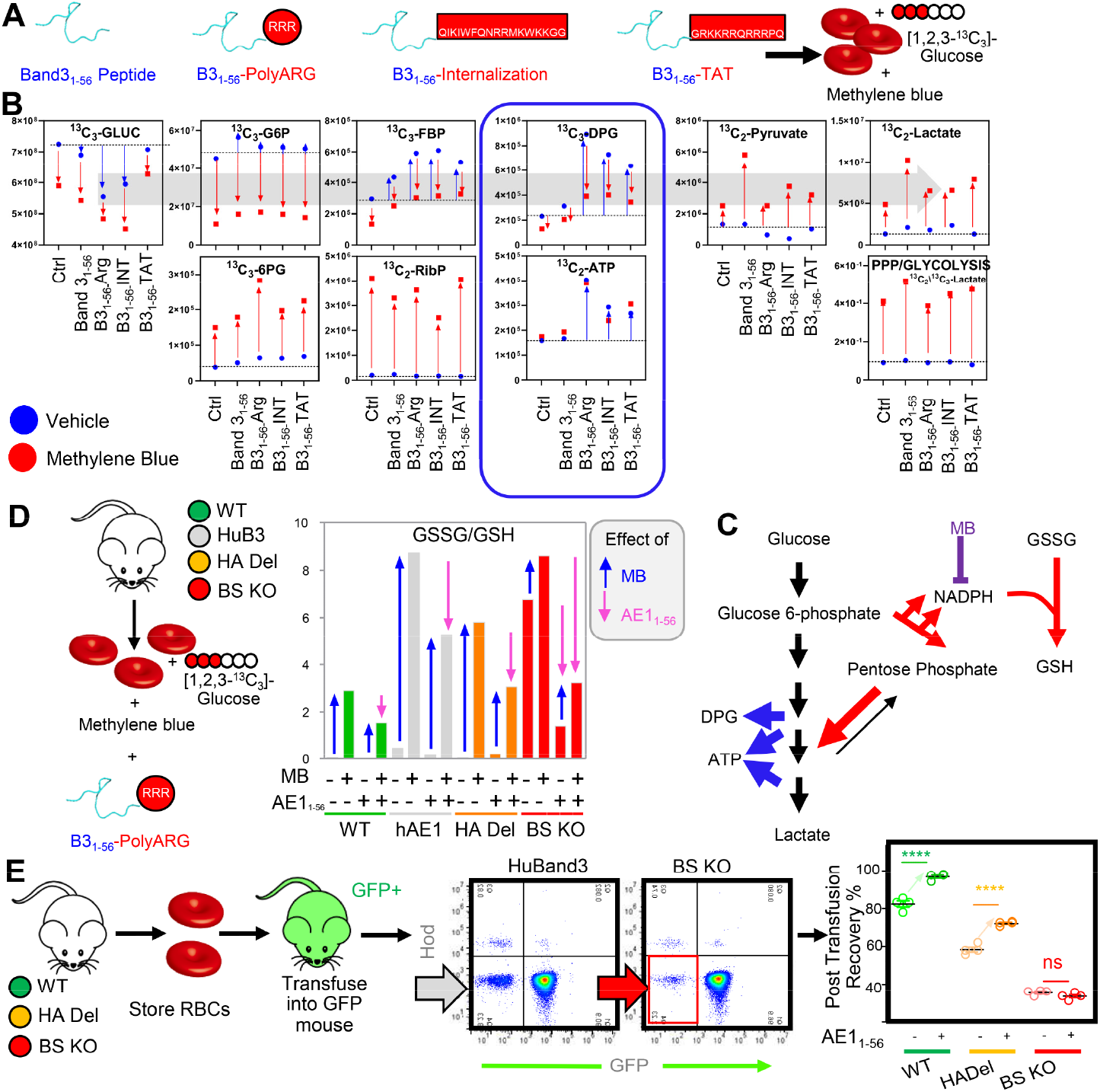
Cell membrane-permeable band 3_1-56_ peptides reverse the defect in the glutathione recycling capacity of RBCs from band 3 KO mice. Three membrane permeable versions of the band 3 (1-56) peptide were generated through addition of a poly arginine (polyArg), internalization sequence or TAT sequence at the C-term of the peptide. Human RBCs were thus incubated with a control (scramble), a band 3 1-56 peptide (non-penetrating) or the three penetrating peptides in presence of 1,2,3-^13^C_3_-glucose and methylene blue stimulation to activate the pentose phosphate pathway (**A**). In **B**, 13C isotopologues of glycolysis and pentose phosphate pathway are reported for all groups at baseline and following methylene blue stimulation of healthy human RBCs, showing increases in PPP activation following MB in all cases, even though ATP levels were only preserved in the RBCs treated with the cell penetrating peptides, suggesting a metabolic reprogramming consistent with the schematic in **C**. In **D**, RBCs from WT, humanized band 3 (HuB3) or band 3 KO mice lacking residues 1-11 (HA Del) or 12-23 (BS KO) were stimulated with MB in presence of a cell penetrating version of the band 3 peptide (polyArg). Results show increased glutathione oxidation in response to MB treatment in all cases, but significant rescue by the band 3 peptide, especially in the HA Del and BS KO groups. Similarly, storage of murine RBCs from the WT and band 3 KO mouse strains in presence of the polyArg cell penetrating band 3 1-56 peptide promoted increases in post-transfusion recoveries in WT and HA Del mice, but not in BS KO mice (**E**).

## Discussion

Here we provide the first direct mechanistic evidence of a role for the N-term of AE1 in regulating RBC storage quality. Using multiple state of the art direct and indirect structural approaches, we provide the first experimental overview of the RBC interactome, with a focus on the N-term of band 3. While The structure of the N-terminus of AE1 beyond residue K56 has been solved through crystallography^49^, AE1_1-56_ is an intrinsically disorder domain that has hitherto eluded structural studies. In the present study, we provide conclusive, direct evidence of the interaction between GAPDH and band 3 by (i) thermal proteome profiling; (ii) pull-down and (iii) cross-linking of recombinantly expressed AE1_1-56_; (iv) ITC, (v) NMR, (vi) cross-linking with two different agents (DSSO and DMTMM) of recombinantly expressed GAPDH and AE1_1-56_ or AE1_1-30_ – the former showing a stronger interaction at least in part explaining by a hotspot of crosslinks between E40 on AE1 and GAPDH. Of note, cross-linking studies have hitherto failed to show a direct proximity of the (many) acidic residues of the AE1 N-terminus with GAPDH K and/or D/E residues, with only one cross-link reported between the two proteins bridging peptides spanning from residues 356—384 of AE1.^7^ Finally, (vii) we modeled GAPDH-AE1_1-56_ structural interactions, as constrained by all the data above. We also provide evidence of direct and indirect interactions of the N-term of AE1 with several other glycolytic enzymes and show that the metabolic defect in glutathione recycling of oxidatively challenged or stored RBCs can be rescued in WT human and genetically ablated mouse RBCs through incubation with a cell-penetrating AE1_1-56_ peptide. Notably, activation of the PPP and post-transfusion recoveries were normalized in band 3 KO RBCs lacking AE residues 1-11 (HA Del) upon incubation with a cell-penetrating AE1_1-56_ peptide. To this end, it is worth stressing that the band 3 KO mice phenocopied G6PD deficient RBCs^31,32^ with respect to the decreased capacity to activate the PPP and ultimately poor post-transfusion recovery.

On top of confirming established interactors by finely mapping the interfaces between AE1 and these molecules (e.g., hemoglobin, ankyrin, spectrin, carbonic anhydrase, glycophorin A), we identified other likely interactors including several components of glutathione synthesis (e.g., GCLC, GSS), glutathione recycling (6PGD), glutathione-dependent lipid peroxidation pathways (GPX4), as well as other antioxidant enzymes (peroxiredoxins 2 and 6, catalase). While PRDX2 had been previously suggested to interact with the N-term of AE1,^47^ here we provide direct evidence of this interaction. Tracing experiments suggested that RBCs lacking the N-term of AE1 suffer from an altered degree of de novo glutathione synthesis, at least in part explained by altered glutaminolysis and transamination. Indeed, GOT – postulated to be present^41^ but never before identified through proteomics in RBCs – was identified as a potential interactor to AE1_1-56_.

Of note, in the plasma samples we also immunoprecipitated high mobility group nucleosome binding protein and histone H2A type 2, very basic proteins with likely affinity for the very acidic N-term of AE1. While speculative at this stage, if confirmed, this interaction would suggest a role for RBC hemolysis in mitigating the endogenous danger signaling associated with the release of these damage associated molecular pattern (DAMP) proteins in response to organ damage or traumatic injury or other DAMP-stimulating event.^50^

In conclusion, here we presented data supporting a central role of the N-term of AE1 in RBC structural and metabolic modulation in oxidatively challenged RBCs in vitro and ex vivo (e.g., blood storage). These data were here accompanied by preliminary evidence of the possibility to rescue altered RBC metabolic phenotypes upon AE1 genetic ablation/storage-induced damage by means of a cell membrane permeable version of a peptide coding for AE1 amino acids 1-56. Further studies will address whether such peptide could rescue oxidant stress-induced lesion to RBC AE1, not just in the context of storage in the blood bank, but also in those pathologies that have been reported to target the N-term of AE1, including COVID-19.^51^

## Acknowledgments

This research was supported by funds from the Boettcher Webb-Waring Investigator Award (ADA), RM1GM131968 (ADA) from the National Institute of General and Medical Sciences, and R01HL146442 (ADA), R01HL149714 (ADA), R01HL148151 (ADA, JCZ), R21HL150032 (ADA), from the National Heart, Lung, and Blood Institute. The authors are grateful to Devin P. Champagne for his contribution at the early stages of this study.

## Authors’ contributions

AD and JCZ designed the study. AI, JR, EZE performed structural studies. AI, ZD, KCH performed cross-linking proteomics studies. AH, JCZ performed mouse studies. AD performed metabolomics analyses; AI, MD, KCH performed proteomics analyses; DR and SP enrolled and characterized the band 3 Neapolis investigated in the human studies. GP, MPB directed the REDS-III RBC Omics study and related GWAS analysis. AI, JCZ and AD analyzed data. AD prepared figures and wrote the first draft of the manuscript. All the authors contributed to the finalization of the manuscript.

## Supplementary information

is available for this paper. Source Data for all the figures and analyses in this study are enclosed within Supplementary Table 1, divided by figures and experiments.

## Supplementary Figures

**Supplementary Figure 1.**
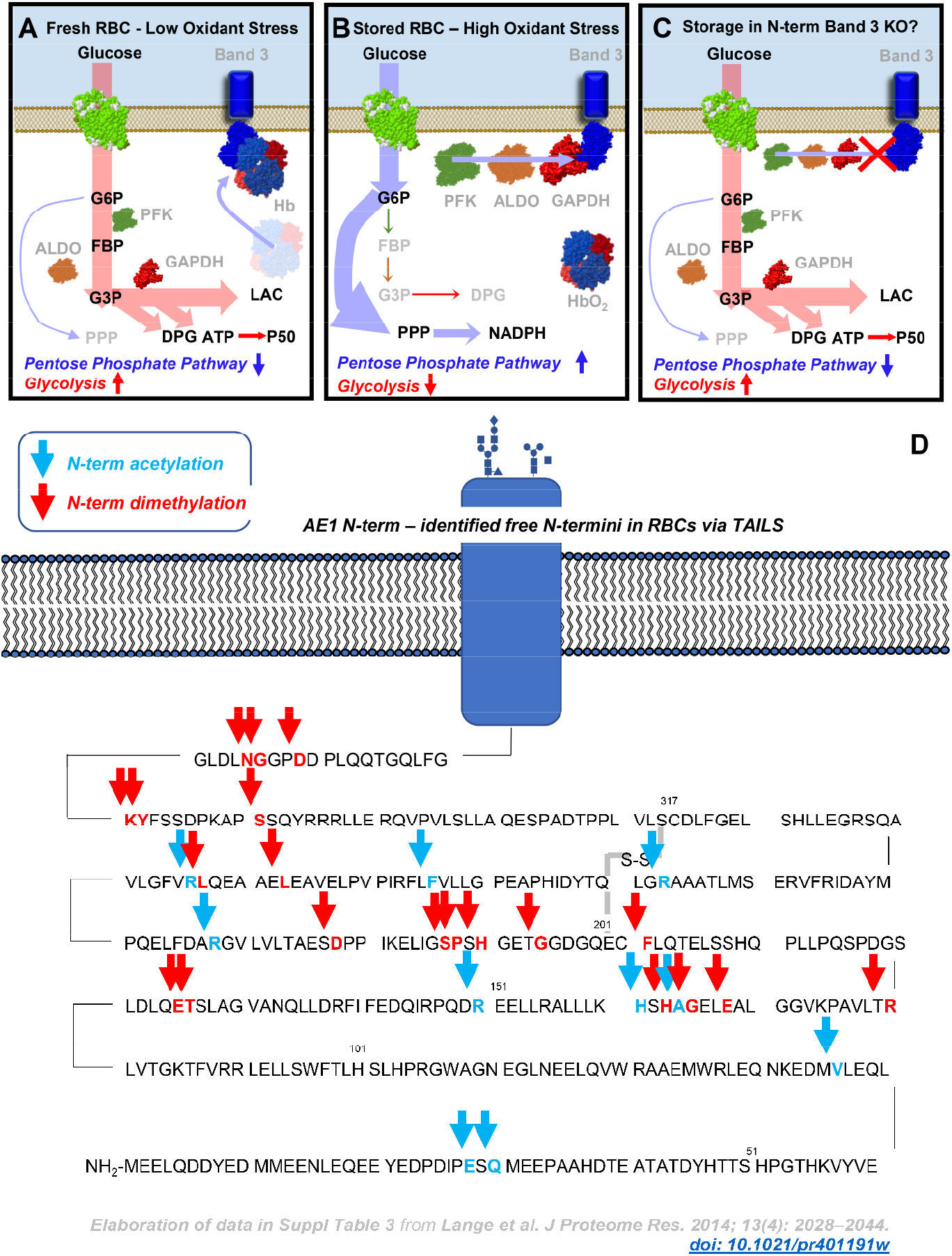
A schematic model of the band 3-dependent regulation of glycolysis and the pentose phosphate pathway, as proposed by Low’s, Castagnola’s, Xia’s and Doctor’s groups. At low oxygen saturation, deoxyhemoglobin binds to the N-term of band 3, while glycolytic enzymes are displaced from the same region and are active – resulting in increased fluxes through glycolysis and decreased fluxes through the pentose phospahte pathway (**A**). At high oxygen saturation, hemoglobin is displaced from the membrane and glycolytic enzymes bind to the N-term of band 3, resulting in their partial inhibition, decreases in fluxes through glycolysis and increased fluxes through the pentose phosphate pathway to generate the reducing equivalent NADPH necessary to counteract oxidant stress, which in turn increases as a function of Fenton chemistry in presence of oxygen (**B**). In **C**, in light of this model and the existing literature from our group and others, we predict that RBCs from mice lacking the N-term of band 3 would suffer from increased oxidant stress during storage under refrigerated conditions. We anticipate that RBCs from these mice would de facto phenocopy G6PD deficiency owing to the incapacity to redirect glucose oxidation fluxes towards the pentose phosphate pathway (**C**). In **D**, Re-elaboration of TAILS data from Lange et al. showing N-term band 3 residues that are fragmented at the end of the shelf-life of human packed red blood cells (storage day 42).

**Supplementary Figure 2.**
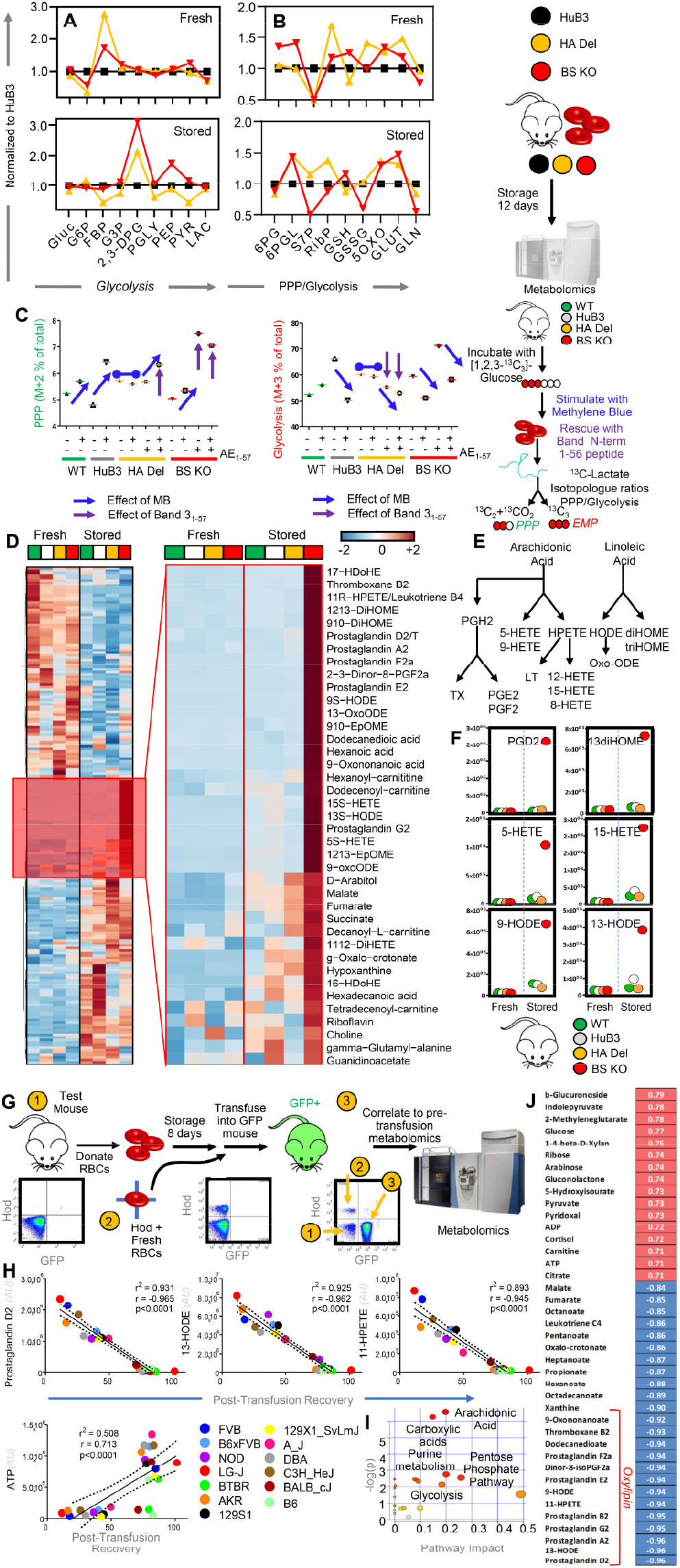
Alterations of glycolysis and the PPP in Band 3 KO mice and metabolic correlates to post-transfusion recovery in mice as gleaned by untargeted metabolomics analyses. Analyses of fresh and stored RBCs from wild type (C57BL6), humanized band 3 mice (HuB3) and mice lacking amino acids 1-11 (HA Del) or 12-23 (BS KO) of band 3 showed strain-specific differences in fresh and stored RBCs in glycolysis (**A**), pentose phosphate pathway (PPP) and glutathione homeostasis (**B**). Data in panels **A-B** report fold-change measurements in HA Del and BS KO mice normalized to the humanized band 3 mice (HuB3). To further investigate the impact on glycolysis and the PPP, RBCs from these four mouse strains were incubated with 1,2,3-13C3-glucose upon stimulation with methylene blue (MB). Rescue experiments were also performed by incubating RBCs with a recombinantly expressed N-term AE1 peptide (residues 1-57 – **C**), prior to determination of lactate isotopologues +3 and +2, deriving from glycolysis and the PPP, respectively (**C**). Metabolomics analyses of stored RBCs from the four strains also showed that RBCs from band 3 KO mice had increased levels of carboxylic acid and lipid peroxidation (**D**), especially metabolites of the arachidonate metabolism (**E**), including prostaglandins, eicosanoids, hydroxy – eicosatetraenoates and octadecenoate (HETEs and HODes – **F**). Metabolomics analyses on fresh and stored RBCs from 13 different mouse strains (**G**) highlight these metabolites as significant correlates to poor posttransfusion recoveries (**H**), as highlighted by the metabolite set enrichment analysis in **I** and the ranked top positive and negative correlates to PTR reported in **J**.

**Supplementary Figure 3.**
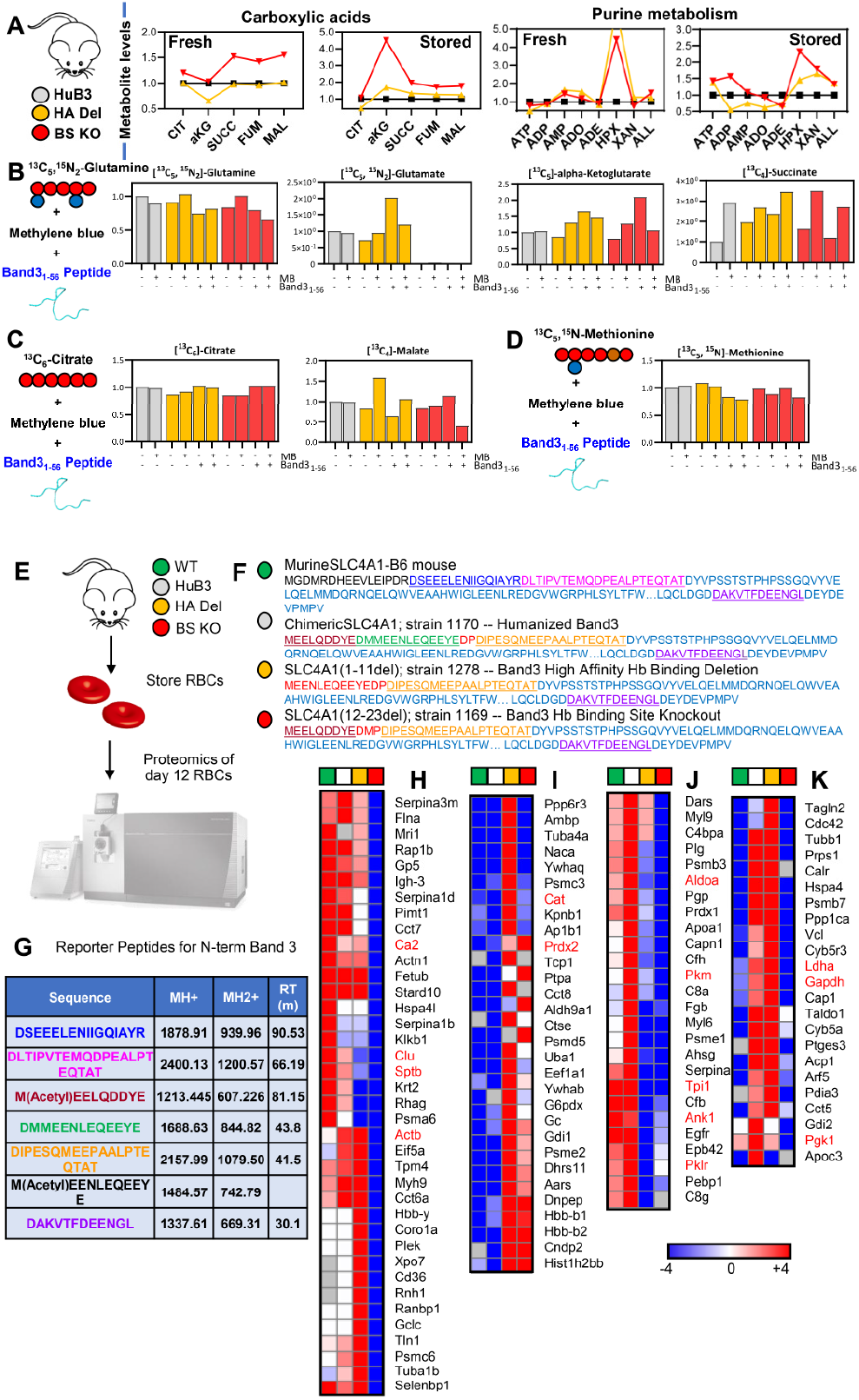
Widespread metabolic alterations were observed in humanized Band 3 (HuB3) or band 3 KO mice lacking residues 1-11 (HA Del) or 12-23 (BS KO). Pathways affected included carboxylic acid and purine metabolism in fresh and stored RBCs (**A**). To further expand on these observations, tracing experiments with ^13^C^15^N-glutamine (**B**), ^13^C-citrate (**C**) and ^13^C^15^N-methionine (**D**) were performed in RBCs from the three mouse strains, in presence or absence of pro-oxidant challenges with methylene blue (MB) and rescue with a recombinant version of the band 3 peptide (residues 1-56). **Proteomics characterization of RBCs from WT and Band 3 KO mice and thermal proteome profiling of recombinant peptide 1-56 of band 3 in RBC lysates.** Proteomics analyses were performed (**E**) to validate the lack of the N-term portion of band 3 in the KO mice (1-11 – highlighted in red – for HA Del; 12-23 – green – for BS KO; in black, peptides unique to the mouse N-term of band 3; in light blue, human band 3 sequence; underlined sequences represent peptides identified in these analyses – **F**). From these analyses, reporter peptide sequences were identified to screen for the four mouse strains through multiple reaction monitoring based approaches (**G**). Proteomics characterization of RBCs from the four mouse strains highlighted a significant impact of BS KO (**H**), either BS KO or HA Del (**I-J**) or HA Del alone (**K**) on several RBC proteins, including numerous well-established interactors of band 3 (highlighted in red in **H-K**).

**Supplementary Figure 4.**
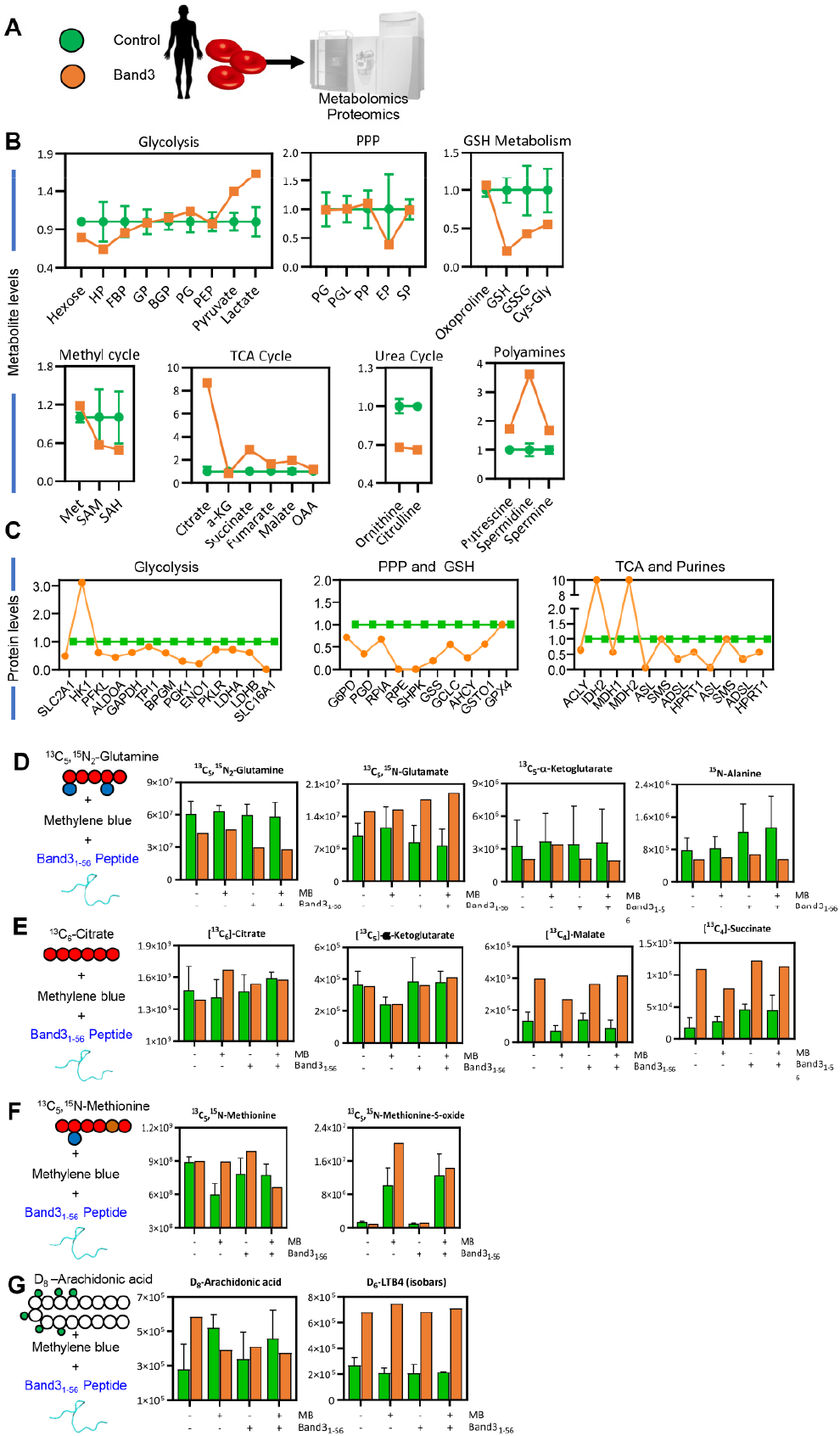
Metabolic impact of band 3 Neapolis in humans. Metabolomics and proteomics analyses were performed on RBCs from a male subject suffering from a genetic mutation resulting in the deletion of the first 11 amino acids of band 3, de facto phenocopying the high affinity deletion KO mice investigated in this study (**A**). Results highlight a significant alteration of glycolysis (increase), glutathione homeostasis (increased oxidant stress) and polyamine metabolism in this subject compared to healthy controls (n=3 – **B**). Proteomics analyses showed higher levels of hexokinase, but lower levels of all the remaining glycolytic enzymes downstream to it, as well as of enzymes of the pentose phosphate pathway and glutathione synthesis in the RBCs from the band 3 Neapolis individual (**C**). Tracing experiments with stable heavy isotope-labeled tracers (^13^C^15^N-glutamine – **D**; ^13^C-citrate – **E**, ^13^C^15^N-methionine – **F**, and deuterium labeled-arachidonate – **G**) confirmed alterations of glutaminolysis, carboxylic acid metabolism, isoaspartyl-damage repair via methylation and oxylipin metabolism (**I**) in the RBCs from the band 3 Neapolis subject – suggesting a role for band 3-dependent regulation of RBC metabolism beyond glycolysis and the pentose phosphate pathway. Specifically, increased consumption of labeled glutamine and generation of labeled glutamate, accompanied by decreases in labeled alphaketoglutarate and alanine are consistent with increased glutaminase activity and decreased transamination in RBCs from the band 3 Neapolis subject compared to controls (**D**). No significant differences were observed with respect to the levels of labeled citrate, or citrate-derived alpha-ketoglutarate (**E**). However, band 3 Neapolis RBCs had four-fold higher levels ^13^C_4_-succinate and malate, in part explained by the higher degree of reticulocytosis in this subject (0.35×106/μl – **Supplementary Table 1**). None of the defects mentioned above was aggravated by treatment with MB or rescue with the recombinant AE1_1-56_ peptide. Studies with labeled methionine showed significant methionine oxidation into its sulfoxide form, especially in response to MB, a defect that was normalized to control values following rescue with the AE1_1-56_ peptide (**F**). Increased arachidonate metabolism in tracing experiments with deuterium-labeled arachidonic acid (**G**), consistent with what observed in band 3 KO mice.

**Supplementary Figure 5.**
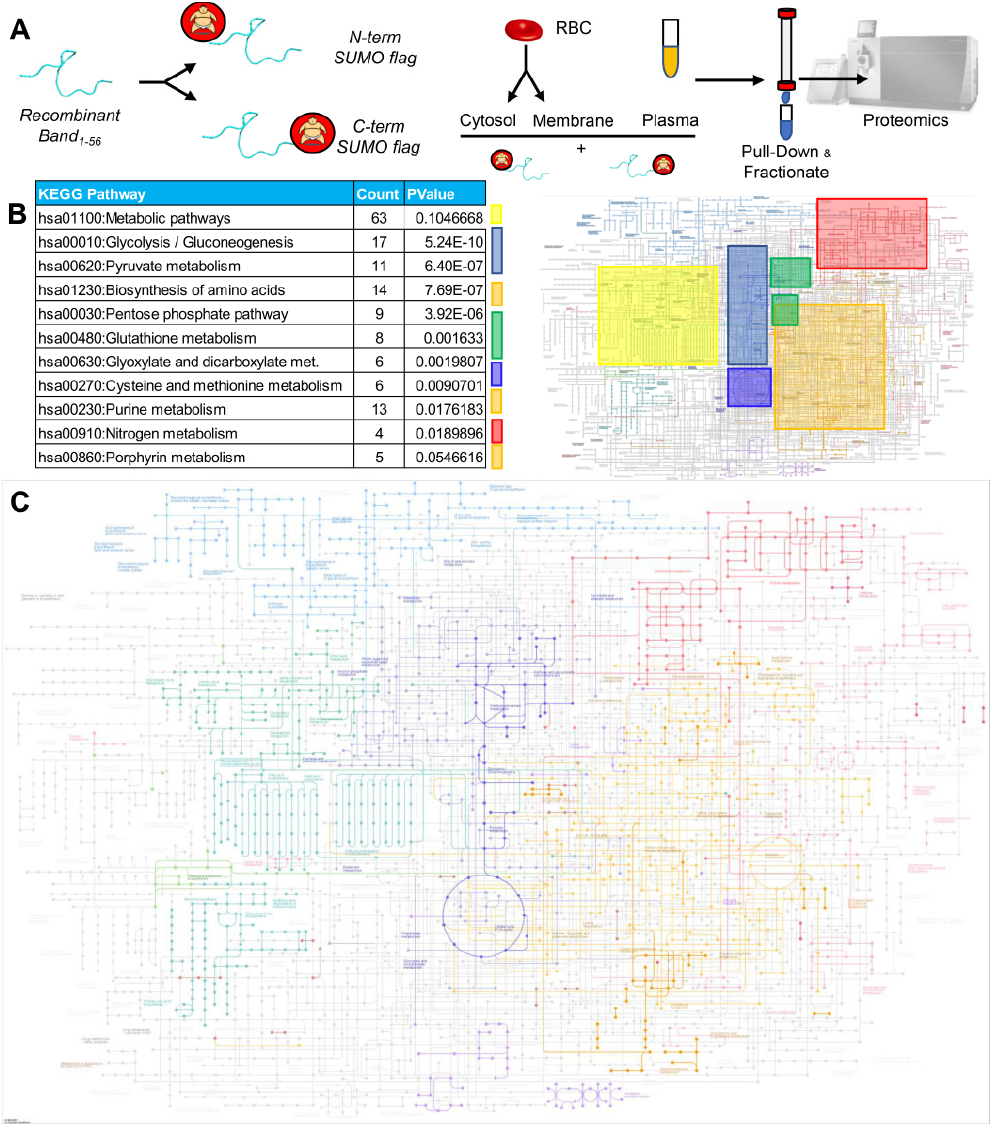
Immuno-precipitation proteomics studies with recombinantly expressed band 3 1-56. A peptide coding for the amino acids 1-56 of the N-term of band 3 was recombinantly expressed with a SUMO-tag at either the N- or C-term terminus of the peptide, prior to incubation with plasma, red blood cell cytosols and membrane in independent experiments, pull-down against the SUMO tag, fractionation and nanoUHPLC-MS/MS-based identification of band 3 interacting partners (**A**). Pathway analyses of the hits from this analysis revealed a widespread interaction of band 3 with up to 63 proteins involved in metabolic regulation, as mapped against the KEGG pathway map of human metabolism (**B**). In **C**, enzymes identified from the pull down of SUMO-tagged band 3 1-56 mapped against the Metabolic pathways map (KEGG map: hsa01100).

**Supplementary Figure 6.**
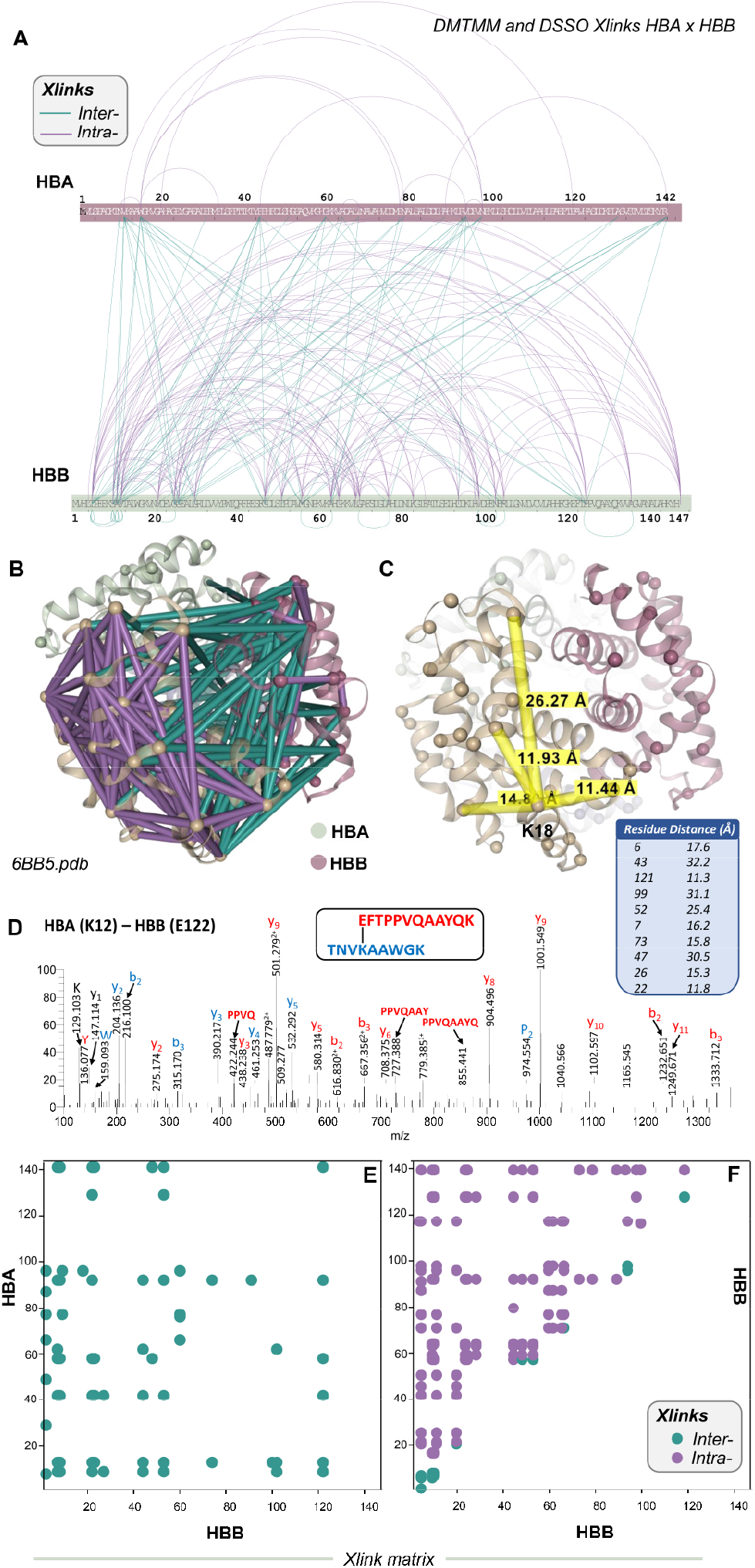
Cross-linking proteomics studies recapitulate structural elucidation of hemoglobin tetramers. In **A**, bar plots show the experimental intra (purple) or inter-molecular crosslinks (teal) between alpha and beta hemoglobin chains (HBA and HBB), respectively. Based on these experimental data, cross-links were mapped against the structure of hemoglobin tetramers for human oxyhemoglobin (6BB5.pdb – **B**). In **C**, highlighted crosslinks and distances between K18 of HBA and neighboring residues. In **D**, a representative mass spectrum from MS3 analyses of one of the highest-scores cross-links, between the peptides containing K12 and E122 of HBA and HBB, respectively (y and b series ions are annotated). In **E** and **F**, matrix plots showing cross-links between HBA and HBB (teal) or within HBB (purple for intra and teal for inter-chain cross-links).

**Supplementary Figure 7.**
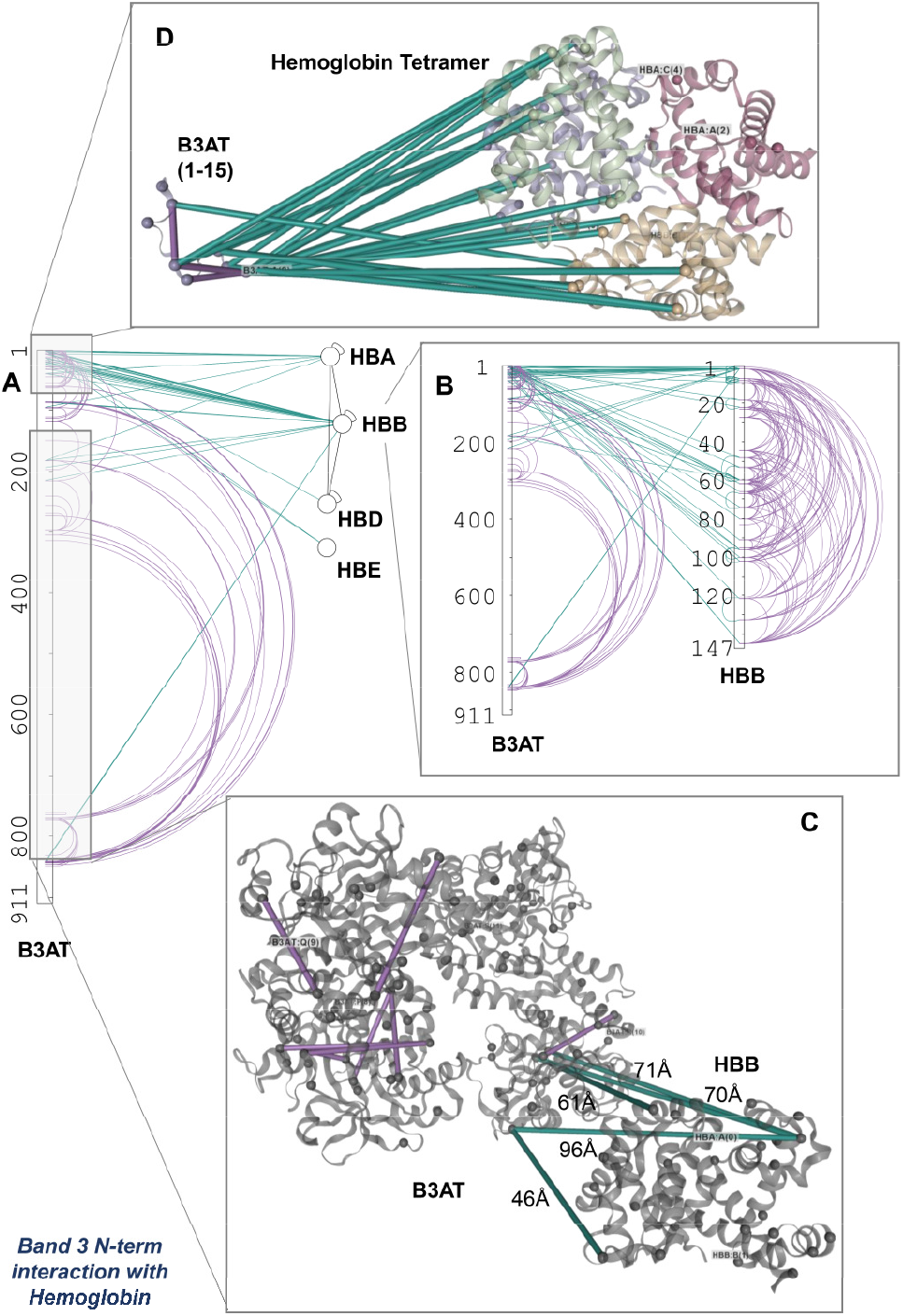
Cross-linking proteomics provides direct evidence of the interactions between hemoglobin and the N-terminus of band 3. These interactions are shown in bar plot format for band 3 (B3AT) with several hemoglobin chains (including alpha, beta, delta and epsilon – HBA, HBB, HBD and HBE – **A**), with a zoom on the cross-links between B3AT and HBB in **B**. In **C** and **D**, the interactions in **B** are highlighted against hemoglobin structure and B3AT cytosolic regions 1-15 and 56-356.

**Supplementary Figure 8.**
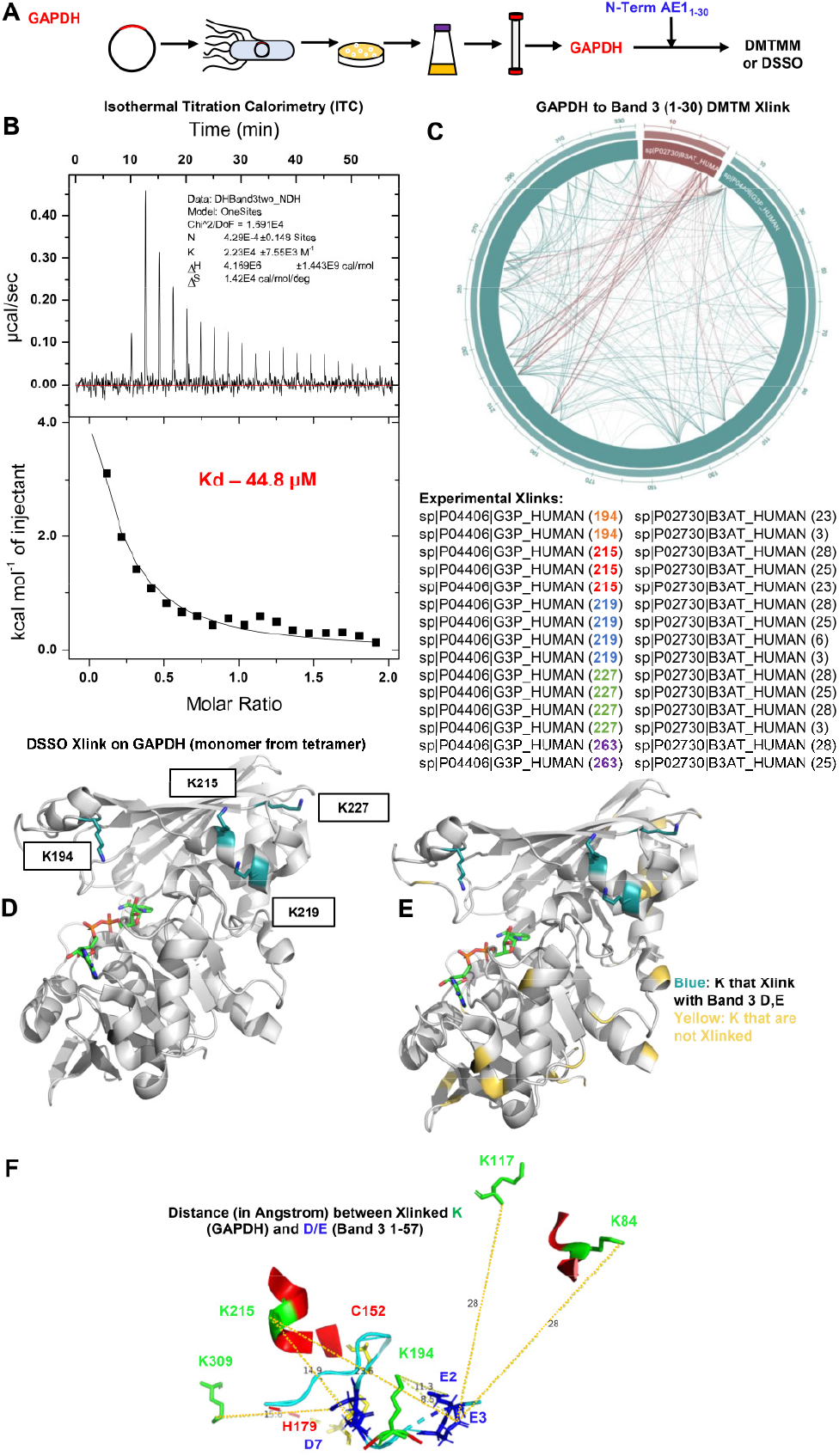
Recombinant expression of band 3 1-30 results in interaction with recombinant GAPDH. **(A)** that have weaker interaction than AE1_1-56_ (44.8 uM vs 2.56 uM). Cross-links with DMTMM are listed in **C** and DSSO cross-links on GAPDH with D/E on band 3 1-30 were mapped against GAPDH structure in **D-E**. In **F**, distance of cross-linked lysine (K – green) *ε*-amine side chain residues on glyceraldehyde 3-phosphate dehydrogenase and carboxylic acid side chains of glutamate/aspartate carboxylate residues on band 3 (residues 1-56 – no K is present on band 3 before residue 57) as experimentally determined by DSSO cross-linking proteomics.

