## Supplementary Methods and Figures for "The interactome of the N-terminus of band 3 regulates red blood cell metabolism and storage quality"

### SUPPLEMENTARY MATERIAL

**SUPPLEMENTARY DATA TABLE** *(PLEAE REFER TO "LEGEND" SHEET WITHIN THE FILE)* .......... **XLSX**

#### Methods

***Animal studies with mice*** All the animal studies described in this manuscript were reviewed and approved by the University of Virginia Institutional Animal Care and Use Committee (protocol n: 4269). Band 3 mouse founders, including huB3, HA Del and BS KO mice – originally generated by Low's group[1] - were acquired from the National Institutes of Health mouse embryo repository and were bred with C57BL/6 females. The use of Ubi-GFP and HOD mice have been previously described in prior work from our group.[2] Whole blood was drawn by cardiac puncture as a terminal procedure for the mice. Blood was snap frozen in liquid nitrogen and storaged at -80ºC until subsequent analysis. For transfusion studies, fresh RBCs (never frozen) were used.

***Band 3 Neapolis*** RBCs (100 ul) were obtained from a subject carrying a rare mutation resulting in the lack of amino acids 1-11 in the N-terminus of band 3, as extensively described by Perrotta et al.[3]

***Methylene blue treatments*** RBCs from WT and band 3 KO mice were incubated with methylene blue (100 uM, Sigma Aldrich) at 37°C for 1h, as described,[4] prior to metabolomics analyses.

***Storage under blood bank conditions*** RBCs from the four main strains investigated in this study, along with 13 different mouse strains described before[5] were collected, processed, stored, transfused and post-transfusion recovery was determined as previously described.[5]

***Tracing experiments with labeled glucose, citrate, glutamine, arachidonic acid and methionine*** RBCs from all the mouse strains investigated in this study (100 ul) or RBC lysates from healthy donor volunteers (n=3) or the individual carrying band 3 Neapolis RBCs were incubated at 37ºC for 1h in presence of 1,2,3-$^{13}C_3$-glucose (5 mM – Cambridge Isotopes – product no. CLM-4673), prior to determination of lactate isotopologues +2/+3 (as markers of pentose phosphate pathway to glycolysis fluxes), as described.[6] Additional tracing experimetns were performed by incubating RBCs with $^{13}C^{15}N$ -methionine (1 mM – Sigma Aldrich – product no. 608106), $^{13}C^{15}N$ -glutamine (1 mM – Cambridge isotope – product no. CNLM-1275-H) or $D_8$-arachidonic acid (Cayman – product no. 390010), as previously described.[7,8]

***Sample processing and metabolite extraction:*** A volume of 50µl of frozen RBC aliquots was extracted in 450µl of methanol:acetonitrile:water (5:3:2, *v/v/v*). After vortexing at 4°C for 30 min, extracts were separated from the protein pellet by centrifugation for 10 min at 10,000g at 4°C and stored at –80°C until analysis.

***Ultra-High-Pressure Liquid Chromatography-Mass Spectrometry metabolomics:*** Analyses were performed using a Vanquish UHPLC coupled online to a Q Exactive mass spectrometer (Thermo Fisher, Bremen, Germany). Samples were analyzed using a 3 minute isocratic condition or a 5, 9 and

17 min gradient as described.[9-11] Solvents were supplemented with 0.1% formic acid for positive mode runs and 1 mM ammonium acetate for negative mode runs. MS acquisition, data analysis and elaboration was performed as described. [9-11]

***Proteomics*** Proteomics analyses were performed via FASP digestion and nanoUHPLC-MS/MS identification (nanoEasy LC 1000 coupled to a QExactive HF, Thermo Fisher), as previously described.[12]

***Recombinant expression of band 3 N-term peptide 1-56***

Unlabeled proteins were grown in Luria broth (LB) while labeled proteins were grown in M9 minimal media (6 g/L $Na_2HPO_4$, 3 g/L $KH_2PO_4$, 0.5 g/L NaCl, 1 g/L $^{15}NH_4Cl$, 2 g/L glucose ($^{13}C_6$-glucose if preparing double labeled protein), 2 mL of 1M $MgSO_4$, 100 µL of 1 M $CaCl_2$, 10 mg/L thiamine) unless stated otherwise.

All Band 3 and GAPDH protein constructs contained an N- or C-terminal 6xHis-SUMO. Constructs were cloned into the bacterial expression vector, pET21b, using the NdeI restriction site. Growth media with proper antibiotics was inoculated with a colony from a freshly transformed LB agar plate and shaken at 37 °C for 16 hours. Fresh media was inoculated with 2.5% of the overnight growth and shaken at 37 °C until an OD of 0.6 was reached. Protein expression was induced by the addition of 1 mM isopropyl-beta-D-thiogalactoside (IPTG). Induced cultures were shaken for 4 hours at 37 °C before being harvested by centrifugation at 4,500 rpm for 10 min.

The purification strategy of all protein constructs was similar. Bacterial pellets were resuspended in Lysis Buffer (50 mM Tris pH 7.5, 300 mM NaCl, 3% glycerol, 10 mM imidazole, 10 mM DTT) and lysed by sonication with 7 cycles of 30 sec on, 30 sec off. Cellular debris was removed by centrifugation at 13,000 rpm for 30 min and 4 °C. Lysate was applied to a column packed with Ni Sepharose Excel resin (Cytiva) and washed with 7 column volumes of Lysis Buffer. Bound protein was eluted with 4 column volumes of Elution Buffer (50 mM Tris pH 7.5, 300 mM NaCl, 400 mM imidazole, 5% glycerol). The elution fractions were pooled and SUMO protease (ulp1, in-house) was added. The sample was dialyzed overnight into Lysis Buffer at 4 °C using dialysis tubing with a 3.5K molecular weight cut-off (MWCO). The sample was applied to a column packed with Ni-NTA His•Bind Resin (Sigma) and the flow-through material was collected. The flow-through fraction was concentrated and applied to a Superose 6 Increase column (Cytiva) equilibrated with Storage Buffer (20 mM Bis-Tris pH 6.5, 150 mM NaCl). Fractions containing target protein were pooled, concentrated, and stored at -80 °C.

***Thermal proteome profiling***

Thermal proteome profiling (TPP) were performed as previously described.[13] Lysates from red blood cells were adjusted to 1 mg/mL and divided into 60 aliquotes of 150 µL. Treated lysate samples were incubated with 100 µM Band 3 peptide prior to heating in a thermo cycler (LifeEco, Bioer) at various temperatures (37, 39, 41.5, 44.9, 49.1, 54.1, 58.5, 61.8, 64.7, 67 °C) for 3 min and immediately cooled to 23 °C. Precipitation was removed by centrifugation at 21,000 rcf for 30 min at 4 °C. A 100 µL volume was taken from each sample and reduced and alkylated (4 M Guanidine HCl, 10 mM DTT, 25 mM IAA) for 30 min in the dark. Four volumes of cold acetone were added to the samples and placed at -20 °C overnight to precipitate protein. Precipitated protein was collected by centrifugation at 18,000 rcf, -5 °C for 25 min. Pellets were washed once with cold acetone and residual acetone was removed with use of a SpeedVac (Thermo Fisher Scientific).

Dried pellets were resuspended in 100 µL of digestion buffer (20 mM triethylammonium bicarbonate (TEAB) pH 8.0, 1 µg Lys-C (in-house), 2 µg sequencing grade trypsin (Promega)). Samples were homogenized with the use of a sonicator (Bioruptor Pico, Diagenode) set at 10 °C for twelve cycles of 30 sec on, 30 sec off. Samples were digested overnight at 37 °C with constant mixing at 800 rpm. Samples were isobaric labeled with TMT reagent (TMT10plex, Thermo Fisher) as recommended by the manufacturer. Briefly, TMT reagent dissolved in 100% anhydrous acetonitrile (ACN) was added to the peptide digests at a final ratio of 4:1 (w:w). Samples were incubated at 23 °C for 1 hour with constant mixing at 800 rpm. The reaction was quenched by adding hydroxylamine to a final concentration of 0.5% and incubating at 23 °C for 30 min. TMT labeled samples were combined and dried using a SpeedVac.

Peptide fractionation was performed on a Gemini NX-C18 column (Phenomenex) using a Dionex UltiMate 3000. Peptides were separated using the following gradient: 3% B (0–16.5 min), 3–45% B (16.5–41.5 min), 45–65% B (41.5–46.5 min), 65–100% B (46.5–48.5 min), 100% B (48.5–53.5 min) with solvent A (10 mM ammonium formate, pH 10.0) and solvent B (10 mM ammonium formate, 75% ACN, pH 10.0) at a flow rate of 0.5 mL/min. Fractions were collected every 0.75 minutes and pooled into groups of three. Pooled fractions were dried down with a SpeedVac and acidified with 0.5% formic acid (FA). Acidified peptides were desalted using C18 Spin Tips (Thermo Fisher Scientific) and stored in 0.1% FA at -80 °C for until further analysis.

TMT labeled peptides were analysed by nano-ultrahigh performance (UHP)LC–MS/MS (Easy-nLC1200, Orbitrap Fusion LumosTribrid, Thermo Fisher Scientific). Sample was loaded directly onto an in-house packed 100 µm i.d. × 250 mm fused silica column packed with CORTECS C18 resin (2.7 µm, spherical solid core). Samples were run at 400 nL/min over a 70-min linear gradient from 4 to 32% acetonitrile with 0.1% formic acid. The mass spectrometer was operated in positive ion data-dependent mode. MS1 scans were run in the orbitrap from 375 to 1,500 m/z at 120,000 resolution. Full scan automatic gain control (AGC) and maximum injection time was set to $4 \times 10^5$ ions and 50 ms. Ions above an intensity threshold of $1 \times 10^4$ were selected for MS2 analysis with fragmentation by collision induced dissociation (CID). Filtering was performed by the quadrupole with an isolation window of 1.2 m/z. AGC and maximum injection time was set to 1 x $10^4$ and 50 ms. Fragmentation was performed by CID with a normalized collision energy of 35%. Detection was set to the ion trap operating in rapid mode. Synchronous precursor selection (SPS) was utilized to co-select 10 MS2 fragments for MS3 analysis. Excluding the precursor, 10 SPS ions were selected within the 400-1200 m/z window. AGC and maximum injection time was set to 1 x $10^5$ and 105 ms. The co-selected SPS ions were fragmented by high-energy collisional dissociation (HCD) with a normalized collision energy of 65%, and analyzed in the orbitrap with a resolution of 50,000 and scan range of 100-500 m/z. SPS-MS3 scan frequency was determined by a 3 s total duty cycle.

Proteome Discoverer 2.2 (Thermo Fisher Scientific) was used for the database search and TMT quantification. Raw data was searched against the human Swiss-prot database. Search parameters included carbamidomethylation-C, TMT 10plex-K, and TMT 10plex-peptide N-terminus as fixed modifications, oxidation-M and pyro-Q N-terminus were set as variable modifications. Up to two missed cleavages were allowed with MS1 and MS2 tolerances set to 20 ppm and 0.5 Da. Reporter ion quantification was set at the MS3 level with an integration tolerance of 0.003 Da. A 65% threshold was set for SPS-MS3 mass match.

Melting curve analysis utilized the lowest temperature condition as the reference. Curve fitting and data normalization was performed as previously described.[14] Briefly, the equation

$$\boldsymbol{f(T) = \frac{1 - plateau}{1 + e^{-(\frac{a}{T} - b)}} + plateau}$$

was used in Prism (GraphPad) for curve fitting and protein $T_m$ determination. The melting point ($T_m$) is described as the temperature at which the melting curve intercepts 50% protein abundance.

***Band 3 Co-IP and chemical cross-linking***

Red blood cell lysates were generated by exposing freshly procured and washed RBCs to hypotonic conditions. Lysates were adjusted with IP Buffer (20 mM HEPES pH 7.4, 100 mM NaCl, 1X Halt Protease Inhibitor Cocktail (Thermo Scientific)). The lysates were centrifuged at 18,000 rpm and 4 °C for 20 min to separate the cytosolic and membrane fractions. The membrane fraction was washed 2 times with IP Buffer. The cytosolic and membrane fractions were diluted to 1 mg/mL with IP Buffer.

Band 3 protein constructs containing either a 6xHis or FLAG tag sequence were used for all co-IP experiments. Cytosolic and membrane fractions were incubated with 50 µM Band 3 protein for 20 min at 4 °C. A second set of samples was prepared for chemical crosslinking with DMTMM and

DSSO (disuccinimidyl sulfoxide, Thermo Fisher) followed by co-IP. In brief, DMTMM and DSSO were used at 20 mM and 1 mM, respectively. The reaction was incubated at 23 °C for 30 min before being quenched with the addition of 40 mM ammonium bicarbonate.

Complexes were precipitated by the addition Ni-NTA (88831, Thermo Fisher) or anti-FLAG (A36797, Thermo Fisher) magnetic agarose. The samples were mixed by rotation for 1 hr at 4 °C. The beads were settled in a magnetic tube rack and unbound material was removed by aspiration. Beads were resuspended with IP Buffer and washed by mixing for 10 min. A total of 6 washes were performed. Bound protein was eluted in 2 rounds with the addition of FLAG peptide (A6002, APExBIO Technology) or 400 mM imidiazole.

The eluted protein was proteolytically digested according to the FASP protocol as previously described[15]. In brief, 300 µg of each sample was reduced, alkylated, and digested at 1:50 with sequencing grade trypsin and mass spectrometry grade rAsp-N (Promega) by incubating at 37 °C for 18 h. Peptides were eluted and acidified to 0.1% formic acid. Crosslinked co-IP samples received an addition enrichment step of strong cation exchange chromatography (SCX) with a Dionex UltiMate 3000 system (Thermo Fisher Scientific). A Proteomix SCX-NP1.7 column (4.6 mm inner diameter, 150 mm length, Sepax Technologies) was used. In brief, peptides were separated using the following gradient: 0% B (0–3.5 min), 0–22.5% B (3.5–18.5 min), 22.5–50% B (18.5–21.5 min), 50–100% B (21.5–23 min), 100% B (23–25.5 min) with solvent A (10 mM KH2PO4, 25% acetonitrile, pH 3.00) and solvent B (10 mM KH2PO4, 25% acetonitrile, 500 mM KCl, pH 3.00) at a flow rate of 0.7 ml/min. Fractions were collected every minute. Fractions 6–26 were pooled into groups of three. All samples were desalted using Pierce C18 Spin Tips (Thermo Fisher Scientific) for subsequent liquid chromatography with tandem mass spectrometry (LC–MS/MS) analysis.

All samples were analysed by nano-ultrahigh performance (UHP)LC–MS/MS (Easy-nLC1200, Orbitrap Fusion LumosTribrid, Thermo Fisher Scientific). Sample was loaded directly onto an in-house packed 100 µm i.d. × 250 mm fused silica column packed with CORTECS C18 resin (2.7 µm, spherical solid core) and run at 400 nL/min over a 70-min gradient from 4 to 32% acetonitrile with 0.1% formic acid. Non-crosslinked and DMTMM treated samples used a MS2 acquisition method. In brief, MS1 scans were run in the orbitrap from 300 to 1,600 m/z at 120,000 resolution. MS2 was performed in a stoichiometric fashion on top ions from each precursor scan and fragmented at a HCD collision energy of 30%. MS2 scan frequency was determined by a 5 s total duty cycle. DSSO treated samples utilized an MS3 acquisition scheme. DSSO treated samples were analyzed as previously described.[16] In brief, MS1 scans were run in the orbitrap from 375 to 1,500 m/z at 60,000 resolution. MS2 was performed in a stoichiometric fashion on top ions from each precursor scan and fragmented at a CID collision energy of 22%. MS2 scan frequency was determined by a 5-s total duty cycle. MS3 was triggered by the targeted mass difference of 31.9721 Da, and was performed as a stepped HCD collision energy of 33 ± 3%. Data acquisition was performed using Xcalibur (version 4.1) software.

Instrument data from the non-crosslinked and DSSO treated samples were directly loaded in to Proteome Discoverer 2.2 and searched against the human Swiss-prot database. Constant search parameters included carbamidomethylation-C as a fixed modification, oxidation-M, and pyro-Q N-terminus as variable modifications, allowing for two missed cleavages. DSSO related variable modifications, DSSO-K, DSSO/amidated-K, and DSSO/hydrolysed-K were added for DSSO specific searches. Precursor mass tolerance was set to 10 p.p.m., with MS/MS mass tolerance set to 20 p.p.m. The XlinkX node was used for DSSO crosslink searches and MS2_MS3 was set for crosslink detection. Results were visualized using xiVIEW.[17] Instrument raw files from DMTMM treated samples were directly loaded into MetaMorpheus and searched using the MetaMorpheusXL task.[18] Search parameters included carbamidomethylation-C as a fixed modification, oxidation-M and pyro-Q N-terminus as variable modifications, allowing for two missed cleavages. DMTMM was added as a selectable crosslinker. Precursor mass tolerance was set to 10 p.p.m., with MS/MS mass tolerance set to 20 p.p.m. Results were visualized using xiVIEW.[17]

***Cross-linking proteomics*** Protein complexes consisting of GAPDH and band 3 truncations were crosslinked with 20 mM DMTMM (4-(4,6-dimethoxy-1,3,5-triazin-2-yl)-4-methyl-morpholinium chloride, Sigma) for 1 hr at room temperature. The reaction was quenched by the addition of ammonium bicarbonate to a final concentration of 50 mM. The crosslinked complex was proteolytically digested according to the FASP (filter-aided sample preparation) protocol as previously described[15]. In brief, ~200 µg of crosslinked sample was reduced, alkylated, and digested at 1:50 with sequencing grade trypsin and mass spectrometry grade rAsp-N (Promega) by incubating at 37 °C for 18 h. Peptides were eluted and acidified to 0.1% formic acid. Enrichment of crosslinked peptides was performed by using strong cation exchange chromatography (SCX) with a Dionex UltiMate 3000 system (Thermo Fisher Scientific). A Proteomix SCX-NP1.7 column (4.6 mm inner diameter, 150 mm length, Sepax Technologies) was used. In brief, peptides were separated using the following gradient: 0% B (0–3.5 min), 0–22.5% B (3.5–18.5 min), 22.5–50% B (18.5–21.5 min), 50–100% B (21.5–23 min), 100% B (23–25.5 min) with solvent A (10 mM KH2PO4, 25% acetonitrile, pH 3.00) and solvent B (10 mM KH2PO4, 25% acetonitrile, 500 mM KCl, pH 3.00) at a flow rate of 0.7 ml/min. Fractions were collected every minute. Fractions 6–26 were pooled into groups of three and desalted using Pierce C18 Spin Tips (Thermo Fisher Scientific) for subsequent liquid chromatography with tandem mass spectrometry (LC–MS/MS) analysis. Crosslinked peptides were analysed by nano-ultrahigh performance (UHP)LC–MS/MS (Easy-nLC1200, Orbitrap Fusion LumosTribrid, Thermo Fisher Scientific). Sample was loaded directly onto an in-house packed 100 µm i.d. × 250 mm fused silica column packed with CORTECS C18 resin (2.7 µm, spherical solid core). Samples were run at 400 nl/min over a 70-min linear gradient from 4 to 32% acetonitrile with 0.1% formic acid. The mass spectrometer was operated in positive ion mode. MS1 scans were run in the orbitrap from 300 to 1,600 m/z at 120,000 resolution. MS2 was performed in a stoichiometric fashion on top ions from each precursor scan and fragmented at a HCD collision energy of 30%. MS2 scan frequency was determined by a 5 s total duty cycle. Data acquisition was performed using Xcalibur (version 4.1) software. Instrument raw files were loaded into MetaMorpheus and searched using the MetaMorpheusXL task.[18] Search parameters included carbamidomethylation-C as a fixed modification and oxidation-M as variable modifications, allowing for two missed cleavages. DMTMM was added as a selectable crosslinker type (-H2O, -18.01056 Da). Residue specificity of DMTMM crosslinks was considered as the following: position 1 to position 2, where position 1 is either the protein N-terminus or Lys and position 2 is either the protein C-terminus, Asp, or Glu. Precursor mass tolerance was set to 10 p.p.m., with MS/MS mass tolerance set to 20 ppm. Results were visualized using xiVIEW[17] and Proxl.[19]

***Isothermal Titration calorimetry (ITC)*** All ITC binding experiments were performed with a MicroCal iTC$_{200}$ (Cytiva) set at 25 °C. GAPDH and Band 3 proteins were dialyzed into matching buffer (20 mM Bis-Tris pH 6.5, 50 mM NaCl, 2 mM DTT). The cell contained GAPDH at 0.1 mM while the syringe contained Band 3 at 1 mM. Reference power was set to 10 µcal/sec with a constant stirring speed of 1000 rpm. A total of 20 injections were performed. The first injection was excluded from data analysis. Experiments were performed in duplicate and the results were analyzed with the Origin ITC module.

***Nuclear Magnetic Resonance and structural models*** $^{15}$N-heteronuclear single quantum coherence (HSQC) spectra were collected at 25ºC for the recombinantly expressed band 3 peptide 1-56 in the presence of 100 and 200 uM GAPDH. Data were collected on a Varian 900 using a standard $^{15}$N-HSQC sequence.

***Statistical Analyses:*** Graphs and statistical analyses (either T-test or repeated measures ANOVA) were prepared with GraphPad Prism 5.0 (GraphPad Software, Inc, La Jolla, CA) and MetaboAnalyst 4.0.[20]

**Supplementary Table 1**

| | Patient | N.V. |
|---|---|---|
| **Red Blood Cells (10^6/µL)** | 3.47 | 4.2-5.5 |
| **Hemoglobin (g/dL)** | 11.3 | 12-18 |
| **Hematocrit (%)** | 36.7 | 37-52 |
| **Mean Corpuscolar Value (fL)** | 105.8 | 80-99 |
| **Mean Corpuscular Hemoglobin (pg)** | 32.6 | 26-31 |
| **Mean Corpuscular Hemoglobin Concentration (g/dL)** | 30.8 | 31-36 |
| **Red Blood Cell Distribution Width – SD (fL)** | 90 | 38-44 |
| **Red Blood Cell Distribution Width – CV (%)** | 24.5 | 11-15 |
| **Platelets (10^3/ µL)** | 659 | 150-450 |
| **Reticulocytes (10^6/µL)** | 0.35 | 0.02-0.1 |
| **Total Bilirubin (mg/dL)** | 10.96 | <1.2 |
| **Indirect bilirubin (mg/dL)** | 10.41 | <0.8 |
| **Lactate dehydrogenase (U/L)** | 867 | 120-240 |
| **Ferritin (ng/mL)** | 103 | 20-400 |

**Supplementary Figures**

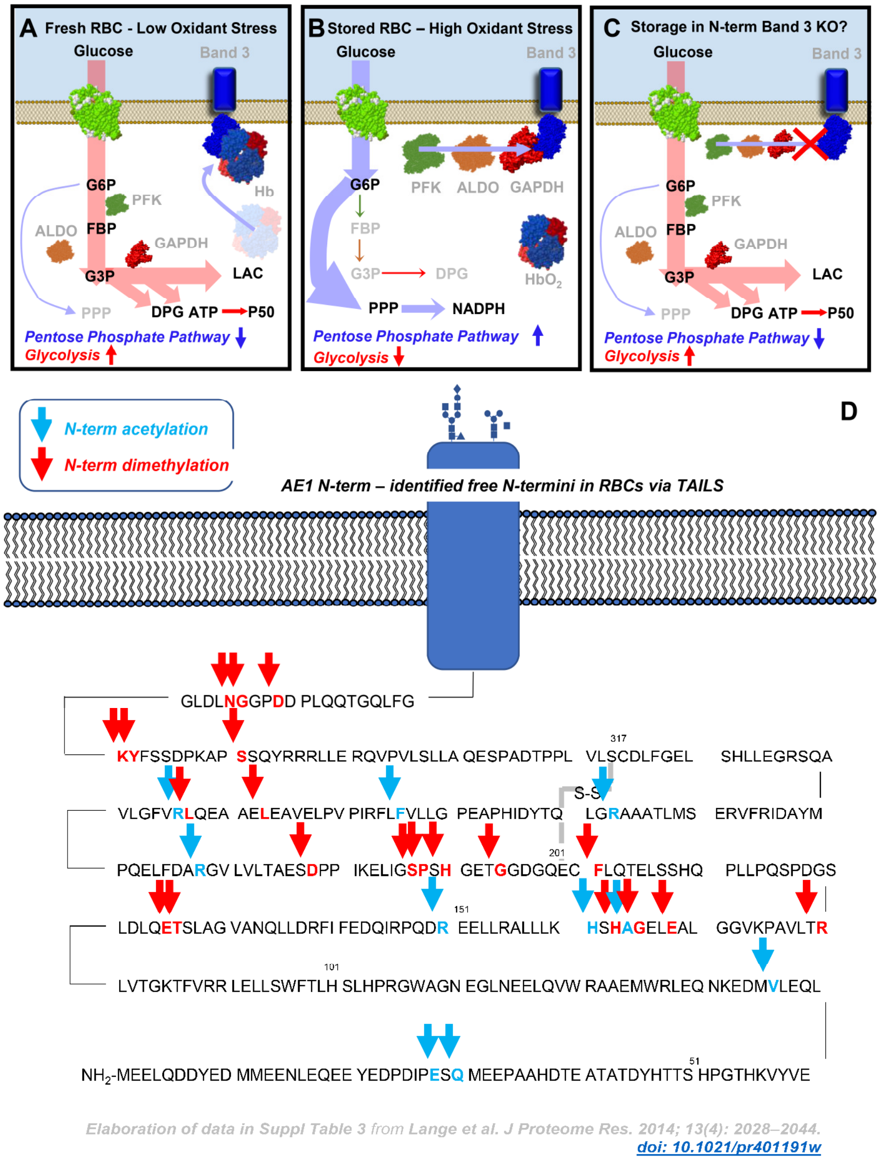

**Supplementary Figure 1 – A schematic model of the band 3-dependent regulation of glycolysis and the pentose phosphate pathway, as proposed by Low's, Castagnola's, Xia's and Doctor's groups.** At low oxygen saturation, deoxyhemoglobin binds to the N-term of band 3, while glycolytic enzymes are displaced from the same region and are active – resulting in increased fluxes through glycolysis and decreased fluxes through the pentose phospahte pathway (**A**). At high oxygen saturation, hemoglobin is displaced from the membrane and glycolytic enzymes bind to the N-term of band 3, resulting in their partial inhibition, decreases in fluxes through glycolysis and increased fluxes through the pentose phosphate pathway to generate the reducing equivalent NADPH necessary to counteract oxidant stress, which in turn increases as a function of Fenton chemistry in presence of oxygen (**B**). In **C**, in light of this model and the existing literature from our group and others, we predict that RBCs from mice lacking the N-term of band 3 would suffer from increased oxidant stress during storage under refrigerated conditions. We anticipate that RBCs from these mice would de facto phenocopy G6PD deficiency owing to the incapacity to redirect glucose oxidation fluxes towards the pentose phosphate pathway (**C**). In **D**, Re-elaboration of TAILS data from Lange et al. showing N-term band 3 residues that are fragmented at the end of the shelf-life of human packed red blood cells (storage day 42).

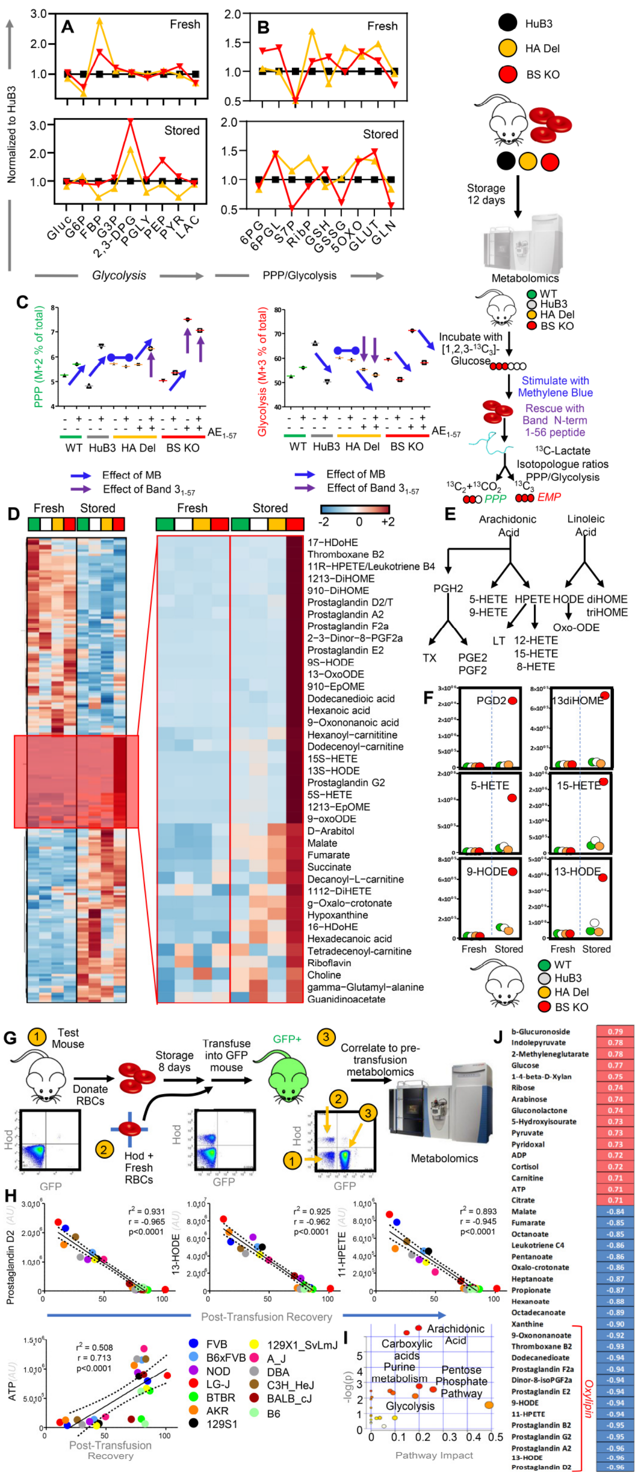

**Supplementary Figure 2 – Alterations of glycolysis and the PPP in Band 3 KO mice and metabolic correlates to post-transfusion recovery in mice as gleaned by untargeted metabolomics analyses.** Analyses of fresh and stored RBCs from wild type (C57BL6), humanized band 3 mice (HuB3) and mice lacking amino acids 1-11 (HA Del) or 12-23 (BS KO) of band 3 showed strain-specific differences in fresh and stored RBCs in glycolysis (**A**), pentose phosphate pathway (PPP) and glutathione homeostasis (**B**). Data in panels **A-B** report fold-change measurements in HA Del and BS KO mice normalized to the humanized band 3 mice (HuB3). To further investigate the impact on glycolysis and the PPP, RBCs from these four mouse strains were incubated with 1,2,3-13C3-glucose upon stimulation with methylene blue (MB). Rescue experiments were also performed by incubating RBCs with a recombinantly expressed N-term AE1 peptide (residues 1-57 - **C**), prior to determination of lactate isotopologues +3 and +2, deriving from glycolysis and the PPP, respectively (**C**). Metabolomics analyses of stored RBCs from the four strains also showed that RBCs from band 3 KO mice had increased levels of carboxylic acid and lipid peroxidation (**D**), especially metabolites of the arachidonate metabolism (**E**), including prostaglandins, eicosanoids, hydroxy – eicosatetraenoates and octadecenoate (HETEs and HODes – **F**). Metabolomics analyses on fresh and stored RBCs from 13 different mouse strains (**G**) highlight these metabolites as significant correlates to poor post-transfusion recoveries (**H**), as highlighted by the metabolite set enrichment analysis in **I** and the ranked top positive and negative correlates to PTR reported in **J**.

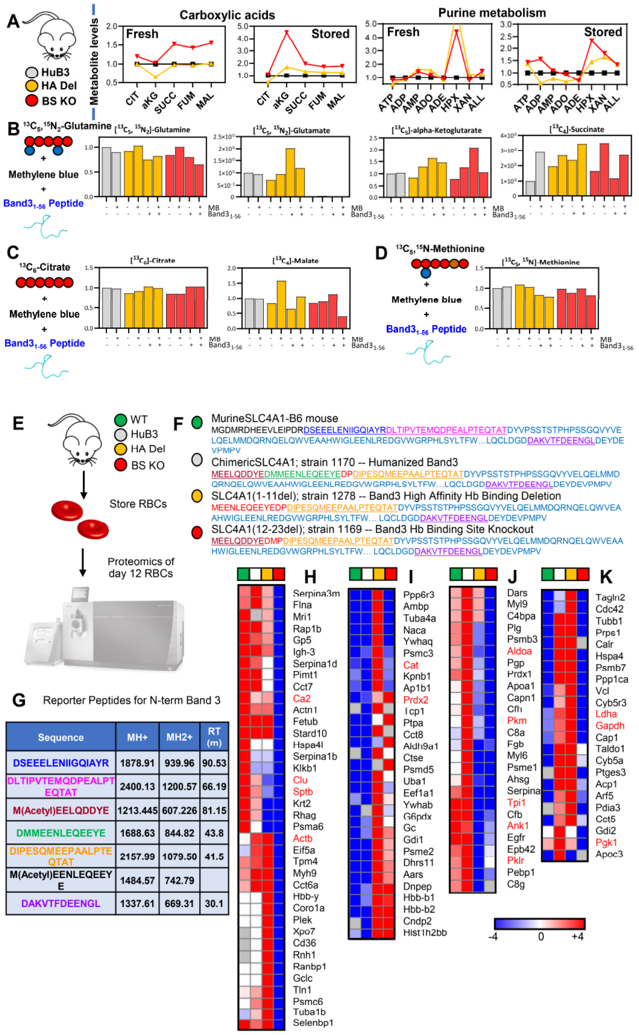

**Supplementary Figure 3 – Widespread metabolic alterations were observed in humanized Band 3 (HuB3) or band 3 KO mice lacking residues 1-11 (HA Del) or 12-23 (BS KO).** Pathways affected included carboxylic acid and purine metabolism in fresh and stored RBCs (**A**). To further expand on these observations, tracing experiments with $^{13}C^{15}N$-glutamine (**B**), $^{13}C$-citrate (**C**) and $^{13}C^{15}N$-methionine (**D**) were performed in RBCs from the three mouse strains, in presence or absence of pro-oxidant challenges with methylene blue (MB) and rescue with a recombinant version of the band 3 peptide (residues 1-56). **Proteomics characterization of RBCs from WT and Band 3 KO mice and thermal proteome profiling of recombinant peptide 1-56 of band 3 in RBC lysates.** Proteomics analyses were performed (**E**) to validate the lack of the N-term portion of band 3 in the KO mice (1-11 – highlighted in red - for HA Del; 12-23 – green – for BS KO; in black, peptides unique to the mouse N-term of band 3; in light blue, human band 3 sequence; underlined sequences represent peptides identified in these analyses – **F**). From these analyses, reporter peptide sequences were identified to screen for the four mouse strains through multiple reaction monitoring based approaches (**G**). Proteomics characterization of RBCs from the four mouse strains highlighted a significant impact of BS KO (**H**), either BS KO or HA Del (**I-J**) or HA Del alone (**K**) on several RBC proteins, including numerous well-established interactors of band 3 (highlighted in red in **H-K**).

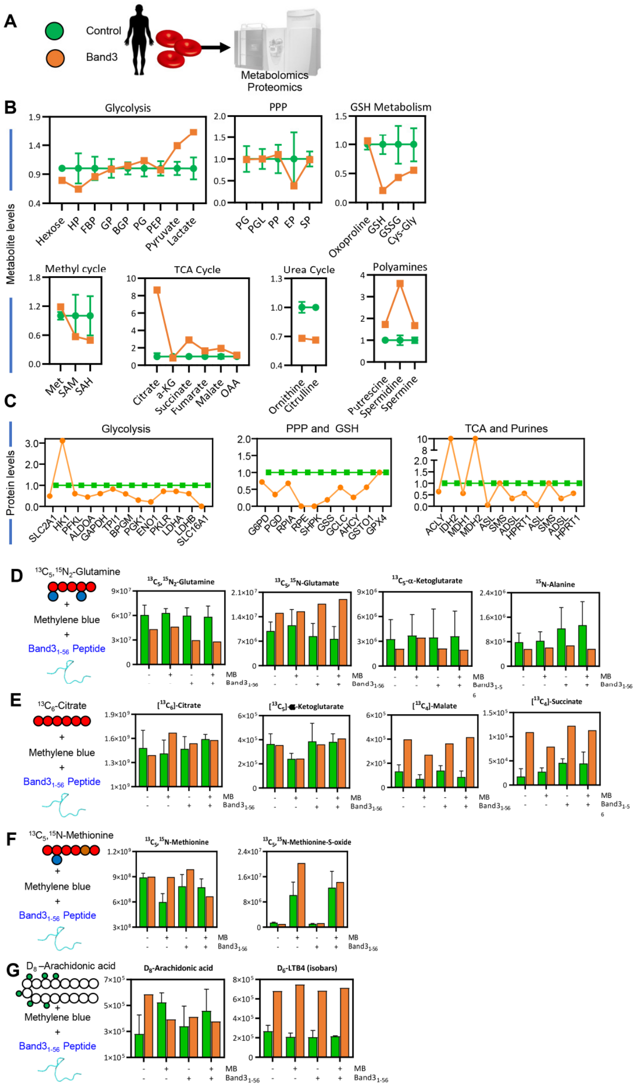

**Supplementary Figure 4 – Metabolic impact of band 3 Neapolis in humans.** Metabolomics and proteomics analyses were performed on RBCs from a male subject suffering from a genetic mutation resulting in the deletion of the first 11 amino acids of band 3, de facto phenocopying the high affinity deletion KO mice investigated in this study (**A**). Results highlight a significant alteration of glycolysis (increase), glutathione homeostasis (increased oxidant stress) and polyamine metabolism in this subject compared to healthy controls (n=3 – **B**). Proteomics analyses showed higher levels of hexokinase, but lower levels of all the remaining glycolytic enzymes downstream to it, as well as of enzymes of the pentose phosphate pathway and glutathione synthesis in the RBCs from the band 3 Neapolis individual (**C**). Tracing experiments with stable heavy isotope-labeled tracers ($^{13}C^{15}N$-glutamine – **D**; $^{13}C$-citrate – **E**, $^{13}C^{15}N$-methionine – **F**, and deuterium labeled-arachidonate – **G**) confirmed alterations of glutaminolysis, carboxylic acid metabolism, isoaspartyl-damage repair via methylation and oxylipin metabolism (**I**) in the RBCs from the band 3 Neapolis subject – suggesting a role for band 3-dependent regulation of RBC metabolism beyond glycolysis and the pentose phosphate pathway. Specifically, increased consumption of labeled glutamine and generation of labeled glutamate, accompanied by decreases in labeled alpha-ketoglutarate and alanine are consistent with increased glutaminase activity and decreased transamination in RBCs from the band 3 Neapolis subject compared to controls (**D**). No significant differences were observed with respect to the levels of labeled citrate, or citrate-derived alpha-ketoglutarate (**E**). However, band 3 Neapolis RBCs had four-fold higher levels $^{13}C_4$-succinate and malate, in part explained by the higher degree of reticulocytosis in this subject (0.35x106/µl – **Supplementary Table 1**). None of the defects mentioned above was aggravated by treatment with MB or rescue with the recombinant AE11-56 peptide. Studies with labeled methionine showed significant methionine oxidation into its sulfoxide form, especially in response to MB, a defect that was normalized to control values following rescue with the AE11-56 peptide (**F**). Increased arachidonate metabolism in tracing experiments with deuterium-labeled arachidonic acid (**G**), consistent with what observed in band 3 KO mice.

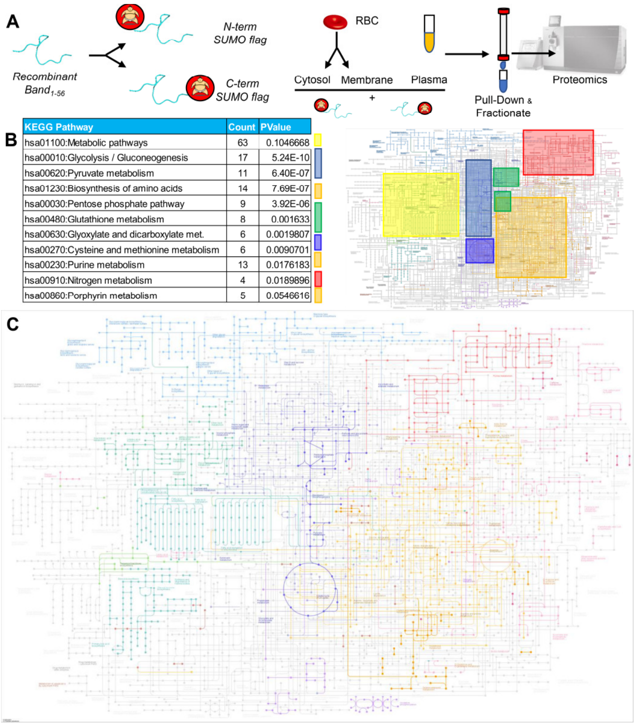

**Supplementary Figure 5 – Immuno-precipitation proteomics studies with recombinantly expressed band 3 1-56.** A peptide coding for the amino acids 1-56 of the N-term of band 3 was recombinantly expressed with a SUMO-tag at either the N- or C-term terminus of the peptide, prior to incubation with plasma, red blood cell cytosols and membrane in independent experiments, pull-down against the SUMO tag, fractionation and nanoUHPLC-MS/MS-based identification of band 3 interacting partners (**A**). Pathway analyses of the hits from this analysis revealed a widespread interaction of band 3 with up to 63 proteins involved in metabolic regulation, as mapped against the KEGG pathway map of human metabolism (**B**). In **C,** enzymes identified from the pull down of SUMO-tagged band 3 1-56 mapped against the Metabolic pathways map (KEGG map: hsa01100).

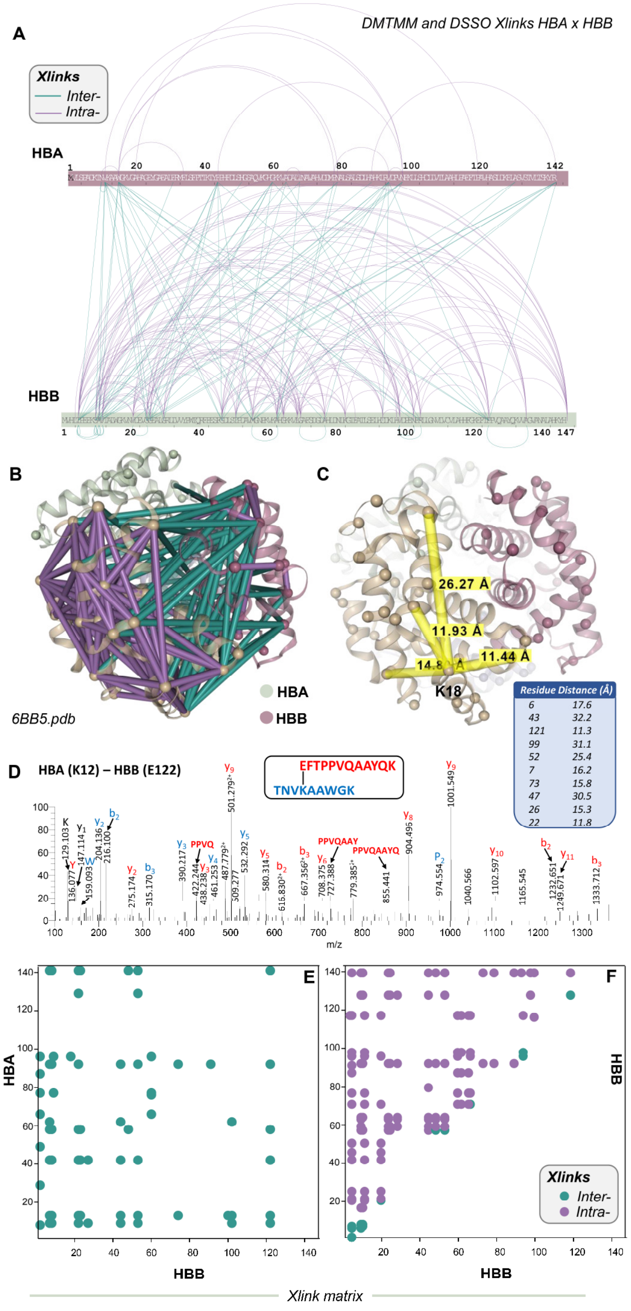

**Supplementary Figure 6 – Cross-linking proteomics studies recapitulate structural elucidation of hemoglobin tetramers.** In **A**, bar plots show the experimental intra (purple) or inter-molecular cross-links (teal) between alpha and beta hemoglobin chains (HBA and HBB), respectively. Based on these experimental data, cross-links were mapped against the structure of hemoglobin tetramers for human oxy-hemoglobin (6BB5.pdb – **B**). In **C**, highlighted cross-links and distances between K18 of HBA and neighboring residues. In **D**, a representative mass spectrum from MS3 analyses of one of the highest-scores cross-links, between the peptides containing K12 and E122 of HBA and HBB, respectively (y and b series ions are annotated). In **E** and **F**, matrix plots showing cross-links between HBA and HBB (teal) or within HBB (purple for intra and teal for inter-chain cross-links).

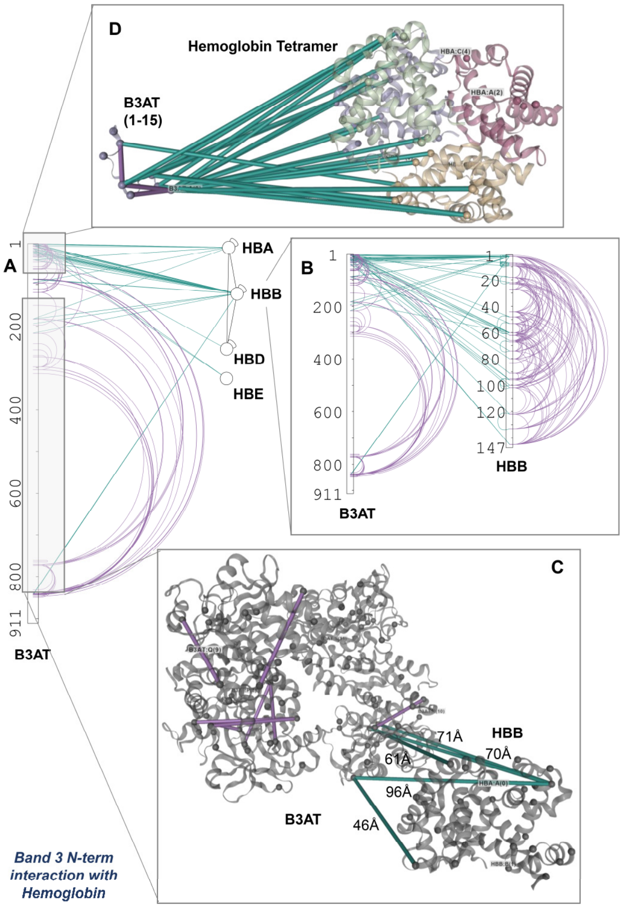

**Supplementary Figure 7 – Cross-linking proteomics provides direct evidence of the interactions between hemoglobin and the N-terminus of band 3.** These interactions are shown in bar plot format for band 3 (B3AT) with several hemoglobin chains (including alpha, beta, delta and epsilon – HBA, HBB, HBD and HBE - **A**), with a zoom on the cross-links between B3AT and HBB in **B**. In **C** and **D**, the interactions in **B** are highlighted against hemoglobin structure and B3AT cytosolic regions 1-15 and 56-356.

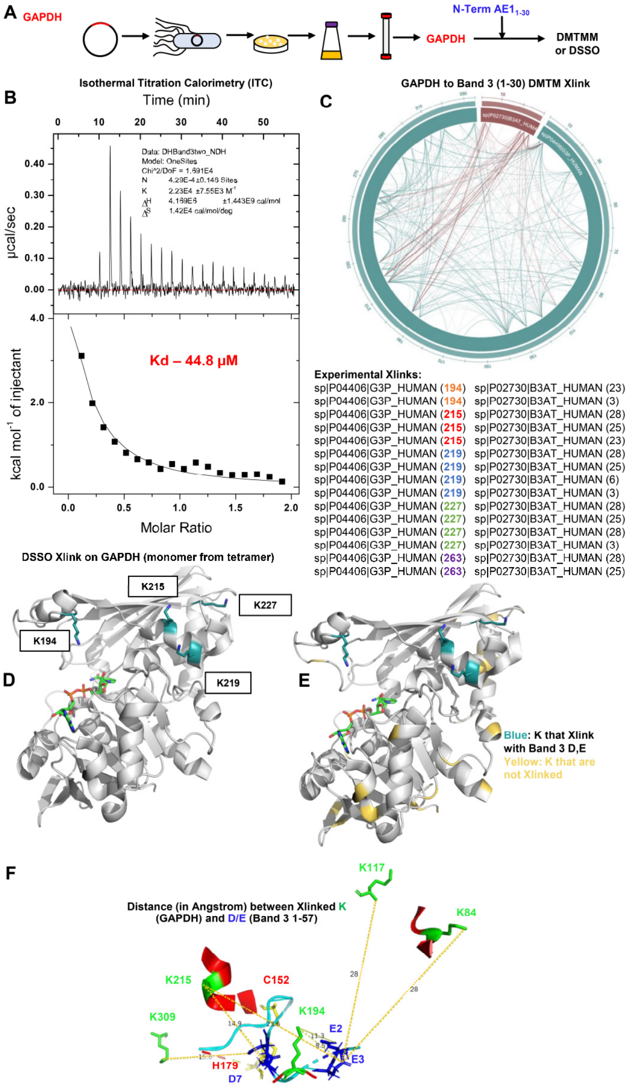

**Supplementary Figure 8 – Recombinant expression of band 3 1-30 results in interaction with recombinant GAPDH (A)** that have weaker interaction than AE1 1-56 (44.8 uM vs 2.56 uM). Cross-links with DMTMM are listed in **C** and DSSO cross-links on GAPDH with D/E on band 3 1-30 were mapped against GAPDH structure in **D-E**. In **F,** distance of cross-linked lysine (K – green) ε-amine side chain residues on glyceraldehyde 3-phosphate dehydrogenase and carboxylic acid side chains of glutamate/aspartate carboxylate residues on band 3 (residues 1-56 – no K is present on band 3 before residue 57) as experimentally determined by DSSO cross-linking proteomics.
